# Predicting ecological interactions across space through pairwise integration of latent network patterns

**DOI:** 10.1101/2025.11.20.689463

**Authors:** Kesem Abramov, Geut Galai, Barry Biton, Rami Puzis, Shai Pilosof

## Abstract

1. Ecological communities are complex and exhibit considerable spatial variability, presenting challenges in accurately understanding these systems. A primary obstacle in ecological research is the existence of ‘missing links’ between species: inevitable unobserved interactions that limit our comprehension of ecological networks and their response to change. While link prediction methods have been developed to address this challenge, most approaches overlook the intrinsic spatial variability of ecological systems.
2. We introduce a flexible, spatially explicit framework based on matrix decomposition that leverages latent structural patterns to predict missing interactions and their strength, without requiring species traits or environmental data. The framework integrates information from paired auxiliary and target networks (locations) using thresholded SVD for link prediction. We applied it to plant–pollinator networks across the Canary Islands, performing pairwise predictions between locations, comparing them to within-location predictions (as a control), and quantifying how spatial variability influences predictive performance.
3. Predictions revealed that latent network structure contains substantial predictive information, with *F*_0.5_ scores consistently exceeding a random baseline (mean *F*_0.5_ = 0.67 ± 0.02 SD), while being less sensitive to interaction strength. The method enabled identifying plausible gaps in the data and producing ecologically coherent predictions. Incorporating information from auxiliary locations enhanced predictive accuracy in certain cases, but success depended on spatial context: predictions were most reliable when derived from nearby, ecologically similar locations, and declined with increasing geographic and ecological distance, consistent with a distance-decay effect.
4. We conclude that the predictability of missing links is spatially variable, reflecting both network and species-level heterogeneity. These patterns provide insights into network structure and the ecological processes shaping it, complementing trait-based approaches. While network structure offers rich predictive information, spatial context is essential for applying it effectively: ignoring spatial variability can obscure ecological signals and inflate predictive error. Our framework is computationally efficient, transferable, and readily applicable to any system with spatial or temporal replication. It can be used for a variety of ecological contexts, including island systems, fragmented landscapes, and environmental gradients, making it a practical and scalable tool for advancing link prediction in ecology.

## 1 Introduction

What we observe in nature is never the full picture. Ecological interaction data are inherently incomplete, limited by sampling effort, methodological biases, and the cryptic nature of many interactions (Chacoff *et al*., 2012; Souza *et al*., 2022; Jordano, 2016; Llopis-Belenguer *et al*., 2023). Unobserved interactions between species pairs fall into two conceptually distinct categories: missing links, which are interactions that occur but went undetected due to sampling limitations, and forbidden links, which cannot occur due to hard biological constraints such as trait mismatches or phenological barriers (Olesen *et al*., 2011). Whether missing or forbidden, all unobserved links between species can obscure key patterns and processes, hindering our understanding of the structure, function, and dynamics of ecological communities (Peralta *et al*., 2024). Given the central role of species interactions in ecosystem functioning and human wellbeing, from crop pollination to disease transmission and biodiversity conservation (Derocles *et al*., 2018; Poisot *et al*., 2023), completing interaction networks can improve predictions of community responses to environmental change (Tylianakis *et al*., 2008; Pearse & Altermatt, 2013). Consequently, there is growing interest in predicting unobserved links (Peralta *et al*., 2024; Desjardins-Proulx *et al*., 2017; Rohr *et al*., 2016; Terry & Lewis, 2020; Strydom *et al*., 2021; Pichler *et al*., 2020; Biton *et al*., 2025). Notably, spatial heterogeneity, a key driver of ecological variability (Poisot *et al*., 2012; Dupont *et al*., 2009), has rarely been explicitly incorporated into link prediction models (Sydenham *et al*., 2022; Dormann *et al*., 2025). Yet incorporating spatial context offers a promising path forward, as it enables the use of variation across networks to inform predictions. Despite this potential, the influence of spatial variability on the accuracy and generalisability of link prediction remains poorly understood.

Spatial variability in interactions is driven by several ecological processes, including changes in species abundance (Vázquez *et al*., 2007), community composition (species turnover) (Vázquez *et al*., 2009), and interaction rewiring (Tylianakis & Morris, 2017; Trøjelsgaard *et al*., 2015). For instance, co-occurring species may interact in one location but not another due to factors such as spatial differences in phenology (Carstensen *et al*., 2014; Olesen *et al*., 2011). Moreover, the degree of interaction turnover varies among species, depending on their tendency to maintain consistent partners across sites (Trøjelsgaard *et al*., 2015; Fortuna *et al*., 2020; Galai *et al*., 2024). These processes often result in distance decay, whereby increasing geographical distance leads to growing dissimilarity in species and interaction composition (Baselga, 2010; Trøjelsgaard *et al*., 2015; Vitali *et al*., 2024; Soininen *et al*., 2007).

Recently, several pioneering attempts have been made to incorporate spatial context into link prediction, aiming to better understand how spatial patterns, such as distance decay, can influence and inform predictive models. Sydenham et al. (2022) used scale-dependent environmental descriptors to predict plant–pollinator interactions in habitat patches. Dormann et al. (2025) integrated abundances, traits, and phylogeny to predict interaction frequencies across replicated networks, revealing that predictability declined with compositional distance. The promising approach of Dansereau et al. (2024) advanced this direction by downscaling a regional metaweb to predict local networks through probabilistic inference, based on species distributions and co-occurrence probabilities. An important consideration arising from these insightful studies is whether data pooling across sites could obscure spatial signals and lead to unrealistic predictions. Such aggregation limits the ability to assess the location-specific informativeness of data. In parallel, several additional studies have shown how information from one network can be “transferred” to inform predictions in other networks (Strydom *et al*., 2023; Caron *et al*., 2024; Nunes Martinez & Mistretta Pires, 2024; Biton *et al*., 2025), considerably advancing the ability to predict interactions in target locations based on data from other areas. However, these methods often rely on traits or phylogeny, which are not always accessible. An alternative approach is to predict links based solely on network structure, i.e., existing linkage patterns between species. Currently, the extent to which structural information from one site can enhance predictions at another within the same spatial system, and how between-site variation in network structure influences predictive success, remain poorly understood.

We introduce a framework that explicitly incorporates spatial variability by predicting interactions and their strength from one site to another, leveraging network structure. Our aim is twofold: (1) to quantify how predictions based solely on local information differ from those that incorporate non-local information, and (2) to identify the processes (e.g., species and interaction turnover) that contribute to predictive success. Specifically, we evaluate whether ecologically meaningful predictions about missing links at a target location can be made using only the structure of interaction networks from an auxiliary location, without relying on trait data. To preserve spatial signals in network structure, predictions are performed in a pairwise framework in which a single auxiliary location is used at a time, rather than pooling information across multiple locations. By focusing on pairwise combinations, we can directly quantify how ecological similarity and spatial separation between locations affect prediction performance, while avoiding the loss of underlying spatial variability in prediction quality that might arise from pooling data across multiple sites. To this end, we develop a spatial prediction pipeline based on singular value decomposition (SVD)—a graph embedding technique that captures latent structural patterns in networks. Recent work has demonstrated the potential of SVD for predicting unobserved links in ecological networks (Strydom *et al*., 2021, 2022; Nunes Martinez & Mistretta Pires, 2024), but it has not been used for spatial analysis. Moreover, prior studies combined it with additional information such as traits or phylogeny (Strydom *et al*., 2022; Nunes Martinez & Mistretta Pires, 2024), which are not always available. In contrast, we rely exclusively on the network structure of the auxiliary and target locations.

We apply our pipeline to an empirical pollination dataset from the Canary Islands, where each island represents a distinct network. This is an ideal model system due to the clear spatial separation between the communities (networks). Because the data set contains weighted links, we are also able to perform both binary and weighted predictions. This is crucial because there have been only a few attempts to predict link weights, with limited success (Terry & Lewis, 2020; Dormann *et al*., 2025). Additionally, because two sites were sampled within each island, we can assess the effect of spatial scale by comparing predictions based on site-level versus island-level (i.e., pooled-site) networks.

We show that predictive accuracy can be enhanced by supplementing focal network data with information from external locations, especially when locations are ecologically or geographically close. Our pipeline produced ecologically coherent predictions that reflected well-documented patterns, despite relying solely on structural information, highlighting the value of accounting for spatial variability for link prediction. To further support exploration of these results, we provide an interactive companion figure at https://ecological-complexity-lab.github.io/svd_based_spatial_prediction/.

## 2 Method description

### 2.1 Data sets used for case studies

#### Main case study

We used a plant-pollinator data set collected by Trøjelsgaard *et al*. (2015) in the Canary Islands, which are volcanic-origin islands in the Atlantic Ocean. A full description of the data set is provided in the original publication. Briefly: the data were collected in five major islands of the archipelago: El Hierro (27.804 N, 17.895 W), La Gomera (28.039 N, 17.226 W), Fuerteventura (28.564 N, 13.891 W), Gran Canaria (27.904 N, 15.433 W) and Tenerife (28.353 N, 16.912 W). Additionally, the data set included a mainland location in Western Sahara (26.161 N, 14.422 W). Owing to its complex geological history, Tenerife was sampled at two distinct locations (Trøjelsgaard *et al*., 2013). Pairwise distances between sampling locations ranged from 52 to 455 km (Trøjelsgaard *et al*., 2015; Vitali *et al*., 2024). At each location, interactions were recorded at two replicate sites situated 50–194 m apart. The sites were selected based on the presence of a large population of the endemic shrub *Euphorbia balsamifera*, and shared additional perennial plant species. Sampling occurred twice between January and March 2010 by conducting 15-minute censuses for every flowering perennial species present. Sites were equally sampled according to accumulation curves and Chao2 estimates (Trøjelsgaard *et al*., 2015). This comparability in sampling completeness allowed us to attribute differences in prediction quality to ecological variation, rather than to inconsistencies in sampling intensity (Vitali *et al*., 2024). The data set contained a total of 249 pollinator species and 39 plant species.

#### Additional data set used to demonstrate generality

To illustrate that the framework is not restricted to spatially replicated pollination networks, we additionally applied the pipeline to a temporally replicated host–parasite interaction dataset. This dataset comprises six weighted interaction networks describing the infection of 22 small mammalian host species by 56 ectoparasite species across six consecutive summers in Siberia (1982–1987) (Krasnov *et al*., 2010; Pilosof *et al*., 2013). Because the objective of this analysis is to demonstrate the applicability of the framework to a different ecological system and type of replication, the results are presented in Appendix S3. In the main text, we discuss, analyse and present results for the primary analysis of the Canary Islands pollination network, unless stated otherwise.

### 2.2 Interaction network construction

Following the original publication (Trøjelsgaard *et al*., 2015), we constructed a plant-pollinator interaction network for each site (14 in total), with rows representing plant species and columns representing pollinator species. In the weighted version of the networks, link weights (matrix cells) were calculated based on flower visitation frequencies and species relative abundances: for each pollinator, a link weight represents the species-specific number of visits paid to a specific plant species multiplied by the relative abundance of the visited plant species (Trøjelsgaard *et al*., 2015). In the binary version, every link whose value is *>* 0 is considered as 1, while a 0 signifies an unobserved interaction. One potential factor that could affect our analysis is the scale at which the data are aggregated (site vs. island). Pooling data across locations is a common strategy to address ecological data sparsity, but it can also introduce spurious interactions (Poisot *et al*., 2015; Blüthgen & Staab, 2024). While the effect of scale on link prediction is a question by itself, which is beyond the scope of this work, pooling data could affect our predictions. Hence, we repeated our analysis at two scales: site and island. For the island scale, we pooled data across the two sites within each island, summing interaction strengths, resulting in 7 island-level networks. We found that the site scale yielded lower quality predictions (Fig. S1), and trends were difficult to detect due to the smaller network sizes and increased noise at this scale, which limited the ability to identify clear relationships. We present the site-level analysis in Appendix S1 (Fig. S1–S7) and proceed throughout the paper with the island scale since this level best addresses the questions we posed and produced better quality predictions.

### 2.3 Link prediction pipeline

#### 2.3.1 SVD algorithm

We inferred missing interactions using a matrix-completion approach based on Singular Value Decomposition (SVD), a widely used dimensionality-reduction technique, which is also commonly applied for link prediction in other fields of network science (Martínez *et al*., 2017). SVD decomposes the observed interaction matrix into a set of latent factors that capture the underlying structure of the network in a reduced space. Specifically, it expresses the original bipartite network as the product of two low-rank matrices, one representing rows (e.g., plants) and the other columns (e.g., pollinators). Each low rank matrix represents species embedded in a shared low-dimensional trait space. Ecologically, the axes of this space can be interpreted as latent interaction traits: unobserved gradients or affinities that influence how species interact. Species with similar interaction profiles are positioned close together along these axes, effectively clustered by their ecological roles or preferences (Strydom *et al*., 2022, 2023).

A standard workflow for link prediction using SVD is to decompose the (possibly partially observed) matrix into its singular vectors and singular values, truncate the decomposition to the top *k* dimensions (latent features), and reconstruct the matrix from these *k* dimensions to obtain predicted values for missing entries (Strydom *et al*., 2023). In a standard SVD workflow, the matrix must be complete (i.e., no NAs); any missing entries (including uncertain non-links, i.e. 0s in ecological networks) must be pre-filled and are then treated as true data in the factorisation. Therefore, a standard SVD cannot exclude selected 0-entries from fitting.

To overcome this limitation, we used the softImpute algorithm (R package softImpute) (Mazumder *et al*., 2010; Hastie, 2022; Hastie *et al*., 2015), which performs regularised matrix completion via iterative soft-thresholded SVD. In contrast to a standard SVD, softImpute works directly with matrices containing missing values (NAs), fitting only to observed entries and iteratively estimating the rest. This makes it possible to withhold both observed links (1s) and non-links (0s) for evaluation. By combining low-rank structure learnt via singular-value optimisation and the ability to exclude both presences and absences during fitting, softImpute avoids biases from arbitrary imputations and handles sparsity naturally, making it particularly well suited for ecological network applications.

softImpute seeks a low-rank approximation *M* that minimises the observed-entry reconstruction error while penalising complexity:

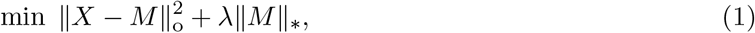

where 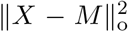 is the sum of squared differences only over observed entries of *X* (the observed network), ∥*M* ∥_∗_ is the nuclear norm (sum of singular values), and *λ* is a nuclear-norm regularisation parameter. At each step, the algorithm iterates between (i) imputing the missing cells with the current low-rank approximation and (ii) applying a soft-thresholded SVD (shrinking small singular values toward zero) to filter out noise, and the process repeats until convergence.

Two parameters control the complexity of the imputed matrix. The first is the maximum rank (*k*), which sets the upper bound on the number of latent features retained in the decomposition. Smaller *k* values produce a simpler model capturing only dominant interaction patterns, whereas larger *k* values allow more nuanced structure but increase the risk of overfitting. We set *k* = 2, as sensitivity checks across a wider grid (*k* = 2, 5, 10) showed no meaningful differences in predictive performance (Fig. S8). The second parameter is the regularisation parameter *λ*, which penalises complexity. Larger *λ* values produce more conservative predictions by applying stronger shrinkage to the singular values, reducing their magnitudes and diminishing the smaller ones most strongly. We set *λ* to the data-derived threshold *λ*_0_ which is the largest value at which the solution does not collapse to an all-zero matrix (Hastie *et al*., 2015; Hastie, 2022). Before applying matrix completion, we centred the interaction matrix, resulting in a matrix with row and column means of zero, to improve numerical stability. After reconstruction, we back-transformed the matrix by restoring the row and column offsets, allowing predicted values to be interpreted on the original scale of interaction strengths.

#### 2.3.2 Matrix aggregation and link withholding

Our goal was to predict interactions in a target location matrix **P** using additional information from an auxiliary location matrix **A**. A special case is **A** = **P**, which yields a self-prediction using only the target network’s own structure (see Experimental design). Performing SVD on **P** and **A** separately would yield different latent spaces, making it impossible to directly infer links in **P** from patterns in **A**. To ensure both matrices are represented in the same latent space, we first combined them into an aggregated matrix **C** (Fig. 1). Aggregating data is a common strategy for representing interactions at broader spatial scales, typically applied to binary presence/absence data (Strydom *et al*., 2023). Since our data contained weighted interactions, we extended this idea to a quantitative setting: for interactions present in both locations (i.e., the same species pair), we summed their weights, while for interactions unique to one location, we retained their recorded values. This fusion preserves both shared and complementary interaction information across locations while providing a consistent basis for decomposition. The resulting matrix **C** can be viewed as a small-scale analogue of the metaweb concept (Dunne, 2006; Poisot *et al*., 2012; Strydom *et al*., 2023), equivalent to a quantitative gamma network encompassing all of the potential interactions occurring in a pair of locations.

**Fig. 1:**
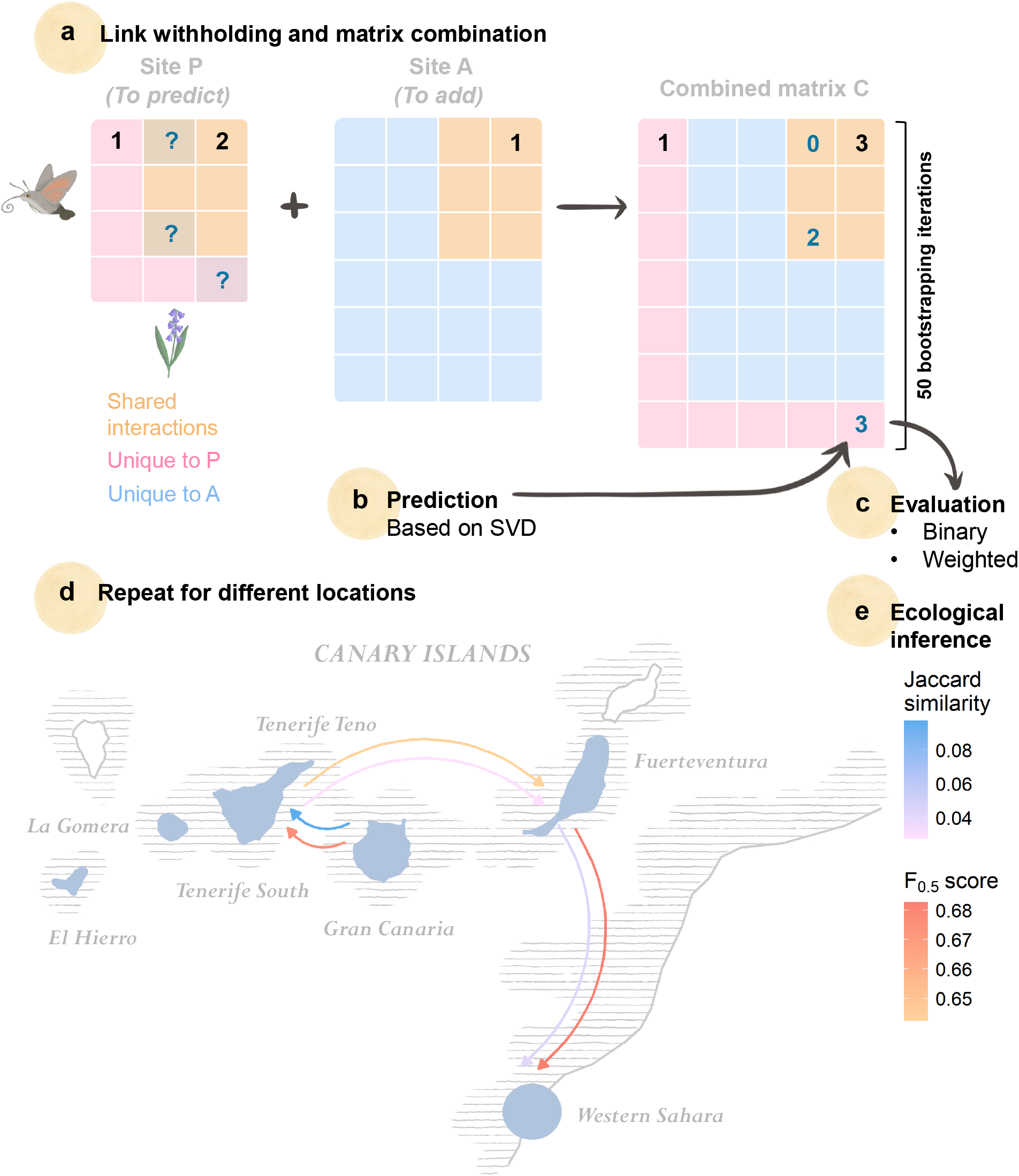
Schematic overview of the spatial link prediction pipeline. **(a)** In each target network (whose links we predict), **P**, we randomly remove 20% of observed links and an equal number of non-links (0s) by setting them to NA. This allows us to simultaneously identify potential missing links and evaluate predictive performance. We then integrate data from an auxiliary location, **A**, to **P**, creating a combined interaction matrix, **C**. To do so, we sum the interaction strength from both locations (represented by the numbers in the matrices). **(b)** SVD is applied to predict the missing values (NA). **(c)** Predictions are evaluated using both binarised and weighted values. **(d)** The entire process is repeated 50 times for each location pair in the Canary Islands. **(e)** For ecological interpretation, we relate predictive performance to potential relevant explanatory variables, including geographical distance and similarity in species composition.

To evaluate predictive performance, we withheld interactions from the target matrix **P** prior to constructing **C**. Specifically, we randomly removed 20% of observed links in **P** by replacing them with NA, analogous to a single fold of 5-fold cross-validation. If a withheld link also occurred in **A**, it was still set to NA in the combined matrix. While this method assumes that a link that was not observed in **P** was also not observed in **A**, it is crucial and necessary to set it to NA to prevent leakage of withheld information from the auxiliary site. This masking is thus a technical evaluation step rather than a biological assumption about whether the interaction occurs at the auxiliary site. Simultaneously, we withheld an equal number of randomly chosen non-links (0s) from **P** by replacing them with NA, and these entries were likewise set to NA in **C**. Thus, both links and non-links selected for evaluation were masked identically in the matrices used for prediction.

Unlike standard link prediction evaluation in ecological applications, this approach evaluates the model’s ability to predict both presences and absences, acknowledging that 0s may reflect incomplete sampling rather than true absences. Using an equal number of links and non-links creates a balanced evaluation set, which allows performance to be compared consistently across different evaluation scenarios because the expected performance of a random predictor remains constant. However, we also tested a strategy that preserves the original class imbalance during withholding. The results of this analysis confirm that predictive performance remains well above the random baseline even under the class imbalance typical of ecological interaction networks (Fig. S9).

While various withholding strategies are possible (e.g., withholding proportionally to species’ degrees), and the choice may influence prediction performance (He *et al*. (2024); see also our case-study demonstration in Fig. S10), we sampled withheld links at random across the matrix to avoid making assumptions about how missing interactions are distributed throughout the network (see Fig. S10 for results under degree-associated withholding strategies).

Importantly, we evaluated predictions only for species present in the target location, even though completion was performed on the combined matrix **C**, which also included species from the auxiliary location **A**. Predictions for **P** were therefore informed by structural patterns in **A**, including interactions involving species absent from the target location. The rationale behind this choice is further explained in the Experimental design.

#### 2.3.3 Model evaluation

We generated predictions from the weighted interaction data and then evaluated the algorithm’s performance in both binary and weighted frameworks. For binary evaluation, raw SVD predictions were transformed to the interval [0,1] using a logistic function (1*/*(1 + *e*^−*x*^)). Each transformed value was then classified as a predicted link (1) or non-link (0) using a probability threshold. We fixed the threshold at 0.7, chosen in preliminary analysis to maximize the *F*_0.5_ score, our primary evaluation metric. *F*_0.5_ is a weighted harmonic mean of precision (the proportion of predicted interactions that are actually true) and recall (the proportion of true interactions correctly predicted; Table S1). Unlike the commonly used *F*_1_ score, which weights precision and recall equally, *F*_0.5_ places greater emphasis on precision. This is particularly appropriate in ecological link prediction, where interaction matrices are sparse and unobserved species pairs are abundant, making false positives (predicted interactions that do not truly occur) a common concern. By giving more weight to precision, *F*_0.5_ provides a more conservative evaluation that penalizes false positives more strongly than false negatives. Higher *F*_0.5_ values indicate that predicted interactions are both accurate and reliable. Because the withheld evaluation set contained equal numbers of interactions and non-interactions (balanced withholding data set), random guessing yields an expected *F*_0.5_ score of 0.5.

In addition to the primary evaluation based on the *F*_0.5_ score, we considered alternative performance metrics to assess the robustness of our results, including the common *F*_1_ score and non-threshold-based evaluators. Specifically, we evaluated predictive performance across classification thresholds by calculating the area under the precision–recall curve (PR-AUC) and the area under the receiver operating characteristic curve (ROC-AUC), which are standard measures of predictive performance across all thresholds.

For evaluating weighted predictions, we first set negative values in the reconstructed matrix to zero, as negative interaction strengths are not ecologically meaningful. This preprocessing step did not affect the selection of the classification threshold or the results of subsequent analyses. We then quantified how closely the predicted interaction strengths (*ŷ*_*i*_) matched the observed strengths (*y*_*i*_) using the normalised Nash–Sutcliffe Efficiency (NNSE) (Dormann *et al*., 2025):

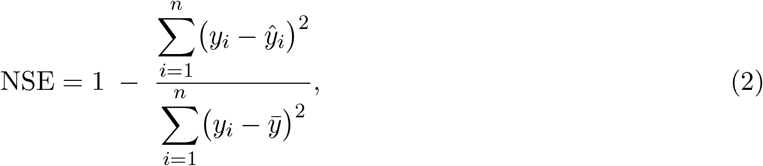

where 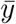 is the mean of the observed values. NSE measures predictive skill relative to simply predicting the mean of the observations. To facilitate comparison across networks, we normalised NSE to the interval [0,1]:

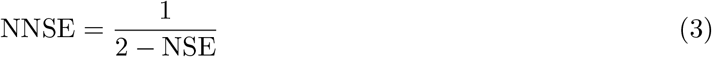

An NNSE of 0.5 indicates no predictive skill (equivalent to predicting 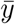), and values closer to 1 indicate higher predictive accuracy. Unlike metrics such as RMSE, NNSE is scale-independent, allowing comparisons between networks with different ranges of interaction strengths.

### 2.4 Experimental design

We applied the prediction–evaluation pipeline to all 42 pairwise combinations of target (**P**) and auxiliary (**A**) island networks, as well as to the 7 cases of self-prediction (target network predicted using only its own data, with no auxiliary location). This yielded 49 prediction cases. For each case, we performed 50 independent iterations of random link withholding, analogous in spirit to repeated cross-validation, to reduce the influence of single withholding configuration on the performance estimates.

While predictions were evaluated only for species present in the target network **P**, the imputation procedure was performed on the combined matrix **C**, which also included species from the auxiliary community **A**. This design allows predictions for **P** to be informed by additional interaction patterns observed in the auxiliary location, providing indirect evidence that can compensate for missing data. At the same time, incorporating species absent from **P** may introduce noise if their interactions in **A** reflect ecological dynamics irrelevant to the target community. To assess this trade-off, we repeated the analysis using only the subset of species shared between the target and auxiliary locations.

### 2.5 Ecological correlates of predictive performance

To explore how predictions vary across space and community structure, we related the cross-location predictive performance to ecological and network-level variables.

#### 2.5.1 Compositional and geographical distance decay

We tested for distance decay in predictive quality by applying a multiple regression on distance matrices (MRM) framework to account for the non-independence of pairwise observations via permutation-based inference (Legendre *et al*., 1994; Lichstein, 2007), following the approach of Vitali *et al*. (2024) applied to the same data set. Analyses were carried out in R using the vegan and ecodist packages (Goslee & Urban, 2007; Oksanen *et al*., 2013).

MRM regresses a symmetric pairwise response matrix against one or more symmetric pairwise predictor matrices. The response matrix contained the mean *F*_0.5_ score for each pair of locations, averaged over the 50 withholding iterations. Because predictions are directional (predicting location *i* using *j* as auxiliary is a distinct case from predicting *j* using *i*), and MRM requires symmetric matrices, we symmetrised the response by averaging the two directed values for each pair: 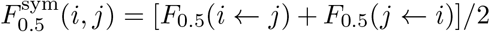. The resulting symmetric 7 *×* 7 matrix was passed to MRM as a lower-triangle distance object, with self-predictions excluded, leaving 21 unique pairs. Predictor matrices of geographic distance, Jaccard similarity, connectance and size are inherently symmetric. Significance was evaluated using permutation tests in which the rows and columns of the response matrix were repeatedly permuted to generate a null distribution of regression statistics, thereby preserving the structure of the predictor matrices while breaking any association with the response.

As geographical distance decay in network similarity arises from species and interaction turnover, we included two ecological predictors: the pairwise geographical distance between locations (km) and the pairwise similarity in interaction composition between networks, measured by the Jaccard index. Apart from these ecological variables, variability in predictive capacity may also partially arise for mathematical reasons: the size (total number of plant and pollinator species) and connectance (proportion of realised links) of the combined matrix **C** are known to correlate with multiple network properties (Delmas *et al*., 2019) and may therefore influence model performance (Farrell *et al*., 2022). Both were therefore included as additional predictors. In total, the MRM predictor set comprised geographic distance, interaction similarity (Jaccard), network size of **C**, and connectance of **C**.

#### 2.5.2 Variable importance

To evaluate which predictors best explained variation in predictive performance, we fitted MRM models for all possible combinations of predictors. Models were ranked using the Akaike Information Criterion corrected for small sample sizes (AICc):

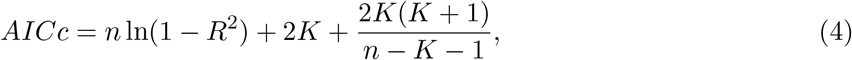

where *n* is the number of pairwise observations and *K* the number of predictors. Because entries in distance matrices are not statistically independent, AICc values were used only for relative model comparison.

We additionally estimated the relative importance of each predictor. Since we perform multiple regressions on pairwise matrices, the conventional methods for calculating relative importance (e.g., using standardized regression coefficients) are not applicable. We therefore estimated the relative importance of each predictor as the reduction in explained variance when that predictor was removed from the full model:

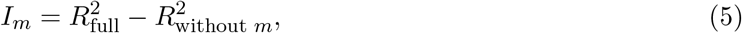

where *I*_*m*_ represents the contribution of predictor *m* to the total variance explained.

## 3 Results

The Canary Islands pollination system consisted of seven networks with sizes ranging from 62 to 84 species and mean connectance of 0.15 ± 0.02 SD (Supplementary Table S2).

Predictive performance was higher when including all species present on both (auxiliary and target) islands, compared to restricting analyses to the subset of species shared between them (Fig. 2a,b). Thus, retaining the full species assemblage across spatial contexts provides more informative predictions. While including all species increases matrix dimensionality and could in principle influence predictive performance simply by increasing matrix size, subsequent analyses indicate that network size itself had little influence on predictive performance (Table 1). Given these findings, we hereafter present results of predictions made with the species of both locations.

**Table 1.** Multiple regression on matrices (MRM) results for models explaining binary predictive performance (*F*_0.5_ scores). The explanatory variables are link composition similarity (Jaccard) between the target and auxiliary networks, geographic distance between networks, and matrix size and connectance of the combined matrix (**C**). The slope coefficients for the four primary explanatory variables are from the full model, quantifying their contributions while controlling for the effects of the other predictors. Relative importance for each model is the reduction in explained variance when that predictor was removed from the full model 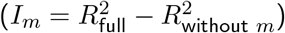.

| Predictor | $R^2$ | $p$-value | AICc | Relative importance |
| --- | --- | --- | --- | --- |
| Distance (km) + Jaccard similarity | 0.429 | 0.005 | -7.094 | 0.299 |
| Jaccard similarity | 0.349 | 0.003 | -6.793 | 0.218 |
| Distance (km) + Connectance + Jaccard similarity | 0.437 | 0.008 | -4.667 | 0.301 |
| Distance (km) + Network size + Jaccard similarity | 0.435 | 0.010 | -4.594 | 0.300 |
| Connectance + Jaccard similarity | 0.352 | 0.020 | -4.429 | 0.220 |
| Network size + Jaccard similarity | 0.349 | 0.008 | -4.338 | 0.218 |
| Distance (km) | 0.176 | 0.050 | -1.852 | 0.081 |
| Connectance + Network size + Jaccard similarity | 0.357 | 0.026 | -1.848 | 0.221 |
| Distance (km) + Connectance + Network size + Jaccard similarity | 0.438 | 0.021 | -1.588 | 0.302 |
| Network size | 0.131 | 0.104 | -0.748 | 0.0002 |
| Distance (km) + Network size | 0.213 | 0.083 | -0.361 | 0.081 |
| Distance (km) + Connectance | 0.187 | 0.128 | 0.313 | 0.083 |
| Connectance | 0.084 | 0.204 | 0.358 | 0.002 |
| Connectance + Network size | 0.132 | 0.261 | 1.686 | 0.002 |
| Distance (km) + Connectance + Network size | 0.219 | 0.173 | 2.208 | 0.084 |

**Fig. 2:**
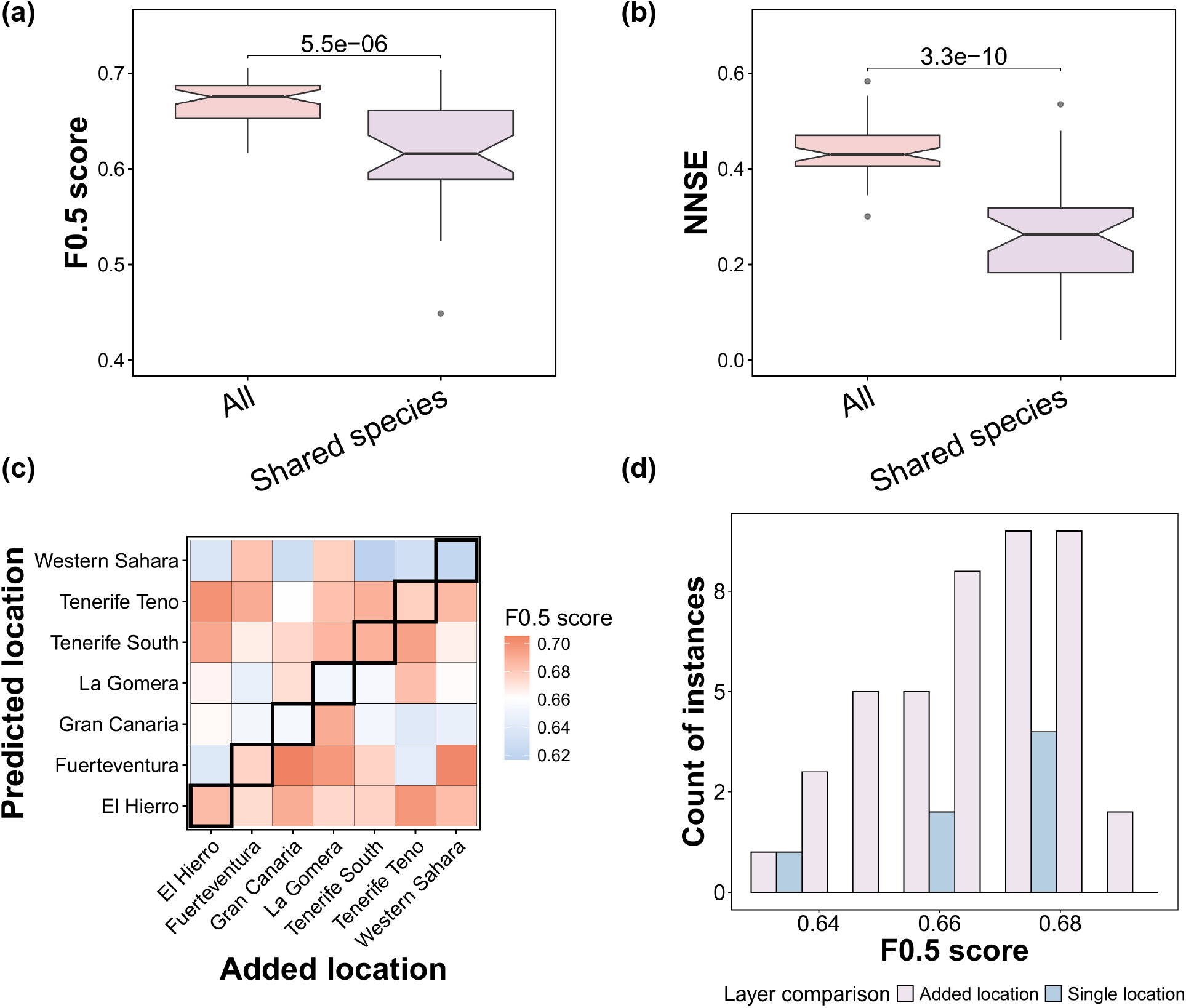
Predictive performance depends on the spatial context. Comparison of predictive performance for binary **(a)** and weighted **(b)** networks using either all species pooled across auxiliary and predicted locations or only overlapping species between locations. P-values correspond to Wilcoxon test for *F*_0.5_ scores and Welch’s test for NNSE. **(c)** Predictive performance of 42 auxiliary-location predictions (off-diagonal values) versus 7 self-predictions (diagonal values, highlighted in a black frame), showing location-specific variability in the predictive power provided by auxiliary information. **(d)** Histogram of values from panel **(c)**, illustrating that the highest *F*_0.5_ scores were generally achieved when incorporating data from auxiliary locations rather than single locations, which also produced large variability in *F*_0.5_.

Overall, the binary predictive performance of the algorithm was strong, with an *F*_0.5_ consistently exceeding 0.5 (the value expected under random guessing) (Fig. 2). The average *F*_0.5_ across the 49 test cases was 0.67 ± 0.02 SD, with a maximum value of 0.71. Results for non-thresholded evaluators followed similar trends, with ROC-AUC and PR-AUC values consistently exceeding random expectations across prediction cases (Fig. S11–S12). In contrast, weighted predictions were not significantly different than random (Fig. 2b), despite a significant correlation between the observed and predicted link weights (Fig. S13). The low predictive power of weights was primarily due to the tendency of the algorithm to assign a wide range of predicted values to the weak observed interactions (Fig. S13).

### 3.1 Evaluating the importance of auxiliary information for predictions

On average, *F*_0.5_ scores were similar between self-predictions (diagonal; 0.66±0.02 SD) and predictions incorporating auxiliary locations (off-diagonal; 0.67 ± 0.02; Fig. 2c,d). However, aggregate statistics mask important location-specific variation. Examining pairwise predictions in detail reveals that in some cases, incorporating data from auxiliary locations produced the highest quality predictions, while in others it reduced accuracy (Fig. 2c,d). Notably, Western Sahara, which was the only mainland location, produced the lowest *F*_0.5_ scores for predictions made using auxiliary island data (Fig. 2c). Together, these results indicate that the benefits of using information from auxiliary locations are location-specific, and depend critically on the ecological compatibility between the target and auxiliary networks.

### 3.2 Interaction-level spatial transferability

Until now, we have analysed predictive power at the location level. Spatial replicability allows us to extend this analysis to variability at the interaction level. We visualised the set of 981 predicted but unobserved links across the Canary Islands pollination system, positioning them alongside the empirically-observed links (Fig. 3a). Most of these interactions (722, 73.6%) were obtained through either self-prediction or auxiliary analyses (Fig. 3b). As expected, these interactions frequently involved the most generalist species, which also occurred on most islands (Fig. 3a, Fig. S14). Nevertheless, 102 interactions (10.4%) of predicted interactions could only be recovered using information from auxiliary locations. Finally, 157 predicted interactions (16%) were recovered exclusively through location-specific information and not with auxiliary data (Fig. 3b). These belonged mainly to species with intermediate or low degree (Fig. 3a), indicating that some interactions can only be detected within their specific spatial contexts.

**Fig. 3:**
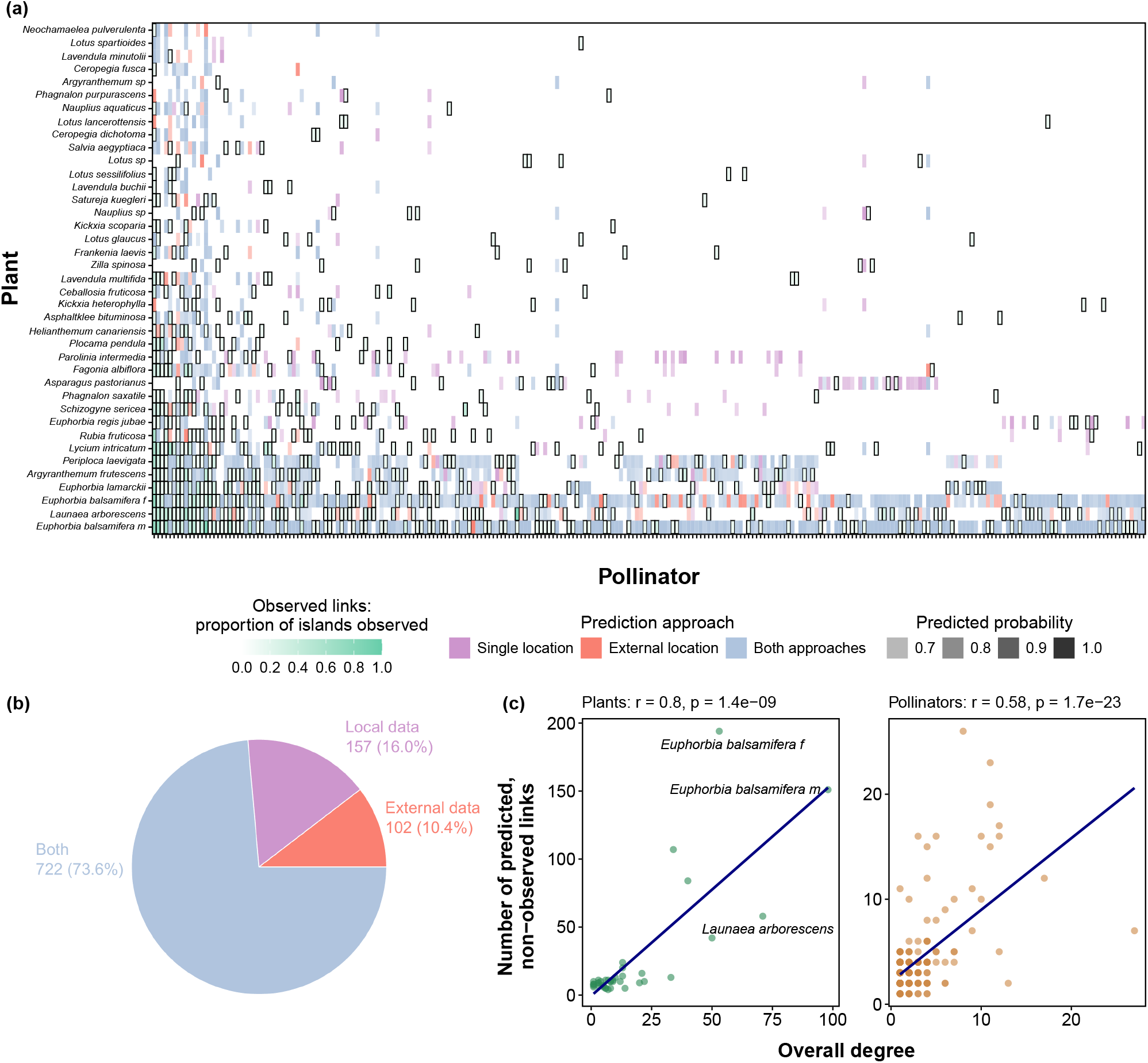
Mapping potential missing interactions in the Canary Islands pollination system. **(a)** Green cells with black borders represent observed interactions; darker shades indicate a higher proportion of islands in which the interaction was recorded. Coloured cells without borders indicate interactions never observed in the data, which were predicted (average predicted probability *>* 0.7 across the 49 prediction cases). The colour denotes the prediction approach, and its intensity reflects the average predicted probability: interactions predicted only when incorporating information from external locations (42 cases) are shown in orange; those predicted using only information from the focal island itself (7 cases) are shown in purple; and those predicted by either approach are shown in blue. Species are ordered according to their overall observed degree (total number of unique interaction partners across islands). **(b)** Amount of unobserved interactions that were predicted using only self-prediction, only using auxiliary islands, or by either of these approaches. The counts correspond to the interactions detailed in **(a). (c)** The number of unobserved yet predicted links per species is strongly correlated with the species’ overall degree across locations (Pearson’s correlation). The two most generalist plants in the system are noted. An interactive version of the figure, with full species labels, information, and threshold controls, is available at https://ecological-complexity-lab.github.io/svd_based_spatial_prediction/.

It is important to note that some predicted links may represent false positives. Typically, validation of such predictions relies on external datasets (e.g., Stock *et al*. (2021); Strydom *et al*. (2022)), but the presence of an interaction in a completely different system does not guarantee its occurrence in the system at hand. Our spatial framework presents an opportunity to overcome this issue because it enables validation within the system itself, thereby increasing confidence in the predicted links. We recalculated the proportions focusing only on the 449 *within-system validated links*, which are the predicted interactions that were observed at least once in the system. Of these, most (397 links, 88%) were recovered by both approaches, 27 (6%) were predicted only from within-island data, and 25 (5.6%) were identified exclusively when incorporating information from other islands.

A noticeable trend was the tendency of the algorithm to assign more missing links to generalists (species with high degree), such as the shrub *E. balsamifera* (Fig. 3a). This was reflected in a strong correlation between the number of predicted links assigned to each species and its total number of observed partners across all islands (Fig. 3c), as well as a clear positive relationship between the local predicted and observed degrees (Fig. S15). Such degree dependence is expected in sparse ecological networks, where species with more observed interactions provide more information for inferring additional links, whereas specialists offer comparatively little (Biton *et al*., 2025). Consistent with this expectation, predictive performance increased with species degree (Fig. S16).

Nonetheless, degree alone did not fully determine prediction patterns. Several highly generalist species, such as *Launaea arborescens*, received only a few additional predicted links (Fig. 3a), despite being consistently among the most connected species across islands (Fig. S17), indicating that this pattern is unlikely to reflect a bias toward local generalists that are not globally generalist. Closer inspection also revealed more nuanced spatial variability (Fig. 3a): missing interactions involving male *E. balsamifera* were consistently recovered regardless of whether local or external data were used, whereas female individuals displayed a different pattern, with certain missing links predicted only when information from other islands was incorporated (Fig. 3a). Finally, the absence of degree dependence in the denser host-parasite network (Fig. S21) suggests that this pattern is system-specific, likely reflecting the sparsity of the pollination network rather than a general property of the algorithm.

### 3.3 Assessing the correlates of spatial predictability

Predictive quality significantly improved with increasing similarity in species composition and interactions between the target and added locations (Fig. 4). We expected that the positive effect of similarity on predictive power would mirror a distance decay in predictive capacity such that the ability to correctly identify links will decline with increasing geographic distance between locations. We indeed detected the expected pattern: although it was marginally significant (MRM, *P* = 0.05; Table 1; Fig. 4d), the geographical distance between locations accounted for 8% of the explained variance in predictive power (Table 1). However, these patterns were not apparent when predicting interaction strength (*P >* 0.14 for correlations of NNSE with geographical distance or species and interaction turnover). Distance decay patterns were also observed in time, when applying the prediction algorithm to a temporal host-parasite network (Fig. S22).

**Fig. 4:**
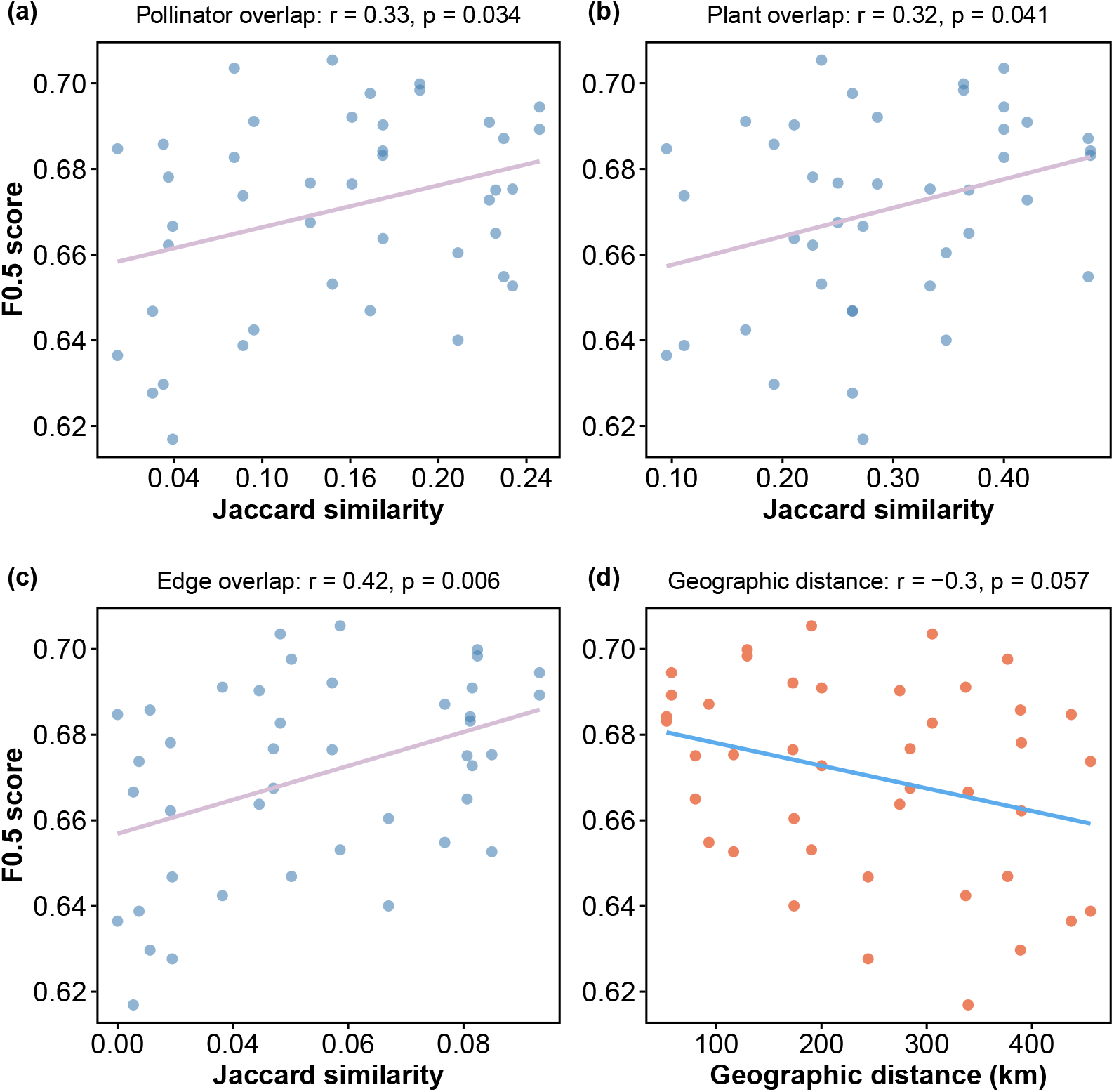
Compositional similarity and interaction consistency enhance predictive accuracy. Predictive ability (*F*_0.5_ scores) increases with greater similarity in pollinator composition **(a)**, plant composition **(b)**, and interactions **(c)** between predicted and added locations. Predictive accuracy declines with increasing geographical distance between predicted and added locations **(d)**. Points represent individual prediction cases (*n* = 42; self-predictions excluded). Reported *R* and *p* values correspond to Pearson’s correlations; see results of permutation-based multiple regression on distance matrices (MRM) in Table 1. An interactive version of the figure, with classification threshold controls and adjustable choice of evaluator, is available at https://ecological-complexity-lab.github.io/svd_based_spatial_prediction/. Across these metrics, the observed trends remain consistent and are more pronounced when using the *F*_1_ score.

An important question is whether predictive ability results solely from ecological variability or also from network properties, which are known to correlate with structure (Delmas *et al*., 2019). To evaluate the joint contribution of these predictors, we fitted a multiple regression on matrices (MRM) including geographic distance, link composition similarity (Jaccard), network size, and connectance (Table 1). The combinations of these predictors yielded 15 competing models. The best-supported model included Jaccard similarity and geographic distance but not matrix size or connectance, indicating that predictive performance is primarily explained by ecological similarity between networks, with an additional contribution of spatial separation. In contrast, matrix size and connectance showed very weak explanatory power and negligible relative importance (Table 1, Fig. S18, Fig. S19). Although network properties did exhibit some effect on performance, these weak effects were primarily noticeable at the site scale (Fig. S6, Fig. S7). As expected, trends in connectance were generally opposite to those observed with network size, with an increase in *F*_0.5_ scores with target matrix size and an increase in NNSE with increasing combined network size and decreasing connectance.

## 4 Discussion

A substantial proportion of ecological interactions remain undetected, even with extensive sampling efforts (Chacoff *et al*., 2012; Jordano, 2016). We developed and tested a spatially explicit framework for link prediction that addresses the long-standing methodological gap of how spatially replicated networks can be leveraged for inference. By leveraging latent structural information from paired auxiliary and target networks, our method accurately predicts missing links and identifies potential data gaps. Our pairwise-location approach represents the opposite end of the spatial aggregation spectrum from the metaweb: rather than aggregating all networks in the system (Strydom *et al*., 2023), we combine only two at a time. While metawebs can be downscaled to local predictions (Dansereau *et al*., 2024) or networks upscaled by aggregation, our method captures the most localized form of cross-site inference.

We found that spatial variability exerts a measurable influence on our capacity to uncover missing interactions, and that integrating external information with focal data can enhance predictive performance. However, predictive ability depends on the ecological compatibility of the added data (e.g., poor prediction of the mainland Western Sahara using additional information from islands). Combining ecologically similar networks yields better predictions, with adjacent locations tending to provide the most compatible data, supporting distance-decay expectations (Soininen *et al*., 2007; Vitali *et al*., 2024; Trøjelsgaard *et al*., 2015). Therefore, aggregating data from dissimilar, distant locations, which is done when predicting links in metawebs (Strydom *et al*., 2023) or multi-site aggregations (Dormann *et al*., 2025), could reduce predictive power.

Our results indicate that network structure encapsulates substantial information for predicting interactions, consistent with previous studies (Terry & Lewis, 2020; Strydom *et al*., 2022, 2023; Biton *et al*., 2025). Degree (the number of unique partners), which reflects a species’ niche breadth, emerged as a key predictor of link assignment in our system. This finding echoes earlier work showing that degree often dominates as a determinant of interaction predictability (Farrell *et al*., 2022; Biton *et al*., 2025). In our spatial context, however, degree also carries a geographic signal, being positively correlated with species occurrence across islands (Fig. S14). Yet degree alone did not fully explain model behaviour: some highly connected species received few additional predicted links (Fig. 3a), the relationship between local observed and predicted degrees was not perfect (Fig. S15), and we also detected intra-specific variation in link assignments (Fig. 3a). Furthermore, degree-dependence in link assignment was not observed in the temporal host-parasite network (Fig. S21).

These patterns suggest that the algorithm captures structural information beyond simple partner counts, integrating subtler spatial and topological signals. Consistent with this, external information uniquely revealed a subset of potential links that are not accessible from local data alone, likely representing spatially variable interactions (Fig. 3b). This highlights the complementary value of auxiliary data: while local network structure is already highly informative for prediction, spatial replication enables the recovery of an additional layer of interactions that would otherwise remain hidden.

Our study shows that spatial information can be leveraged for predicting interactions, yet three main aspects should be further considered in future developments of this approach. First, as with other link-prediction methods, our framework may predict links that do not truly occur in nature. False positive predictions may represent either truly missing interactions or interactions that do not occur — with forbidden links, arising from hard biological constraints such as trait mismatches or phenological barriers, being a subset of the latter. Distinguishing between these cases is a fundamental challenge in ecological link prediction that cannot be resolved without independent empirical corroboration. We partially mitigate this concern by adopting *F*_0.5_ as our primary evaluation metric, which penalizes false positives more strongly than false negatives. A further advantage of our spatial framework is that replication across locations enables *within-system* validation, drawing on the same ecological context rather than relying on external datasets that may differ in community composition or sampling design (Stock *et al*., 2021; Farrell *et al*., 2022). In our analysis, only 31% of the predicted links were observed at least once in the system. Given the limited spatial and temporal coverage of our dataset, some non-validated predictions may nonetheless represent ecologically valid but undetected interactions. False positives can be further reduced by integrating trait-based or phylogenetic information (Peralta *et al*., 2024), or, when extending the method to systems with strong biological constraints — such as seed dispersal or frugivore networks — by incorporating trait or phenology-based masks to exclude biologically impossible species pairs from the candidate interaction set.

Secondly, while binary link predictions were consistently accurate and outperformed random expectations, predictions of interaction strengths performed no better than random and showed limited associations with ecological variables beyond network size and connectance. The low predictive ability of quantitative interactions aligns with previous studies highlighting the difficulty of modelling interaction weights (Terry & Lewis, 2020; Dormann *et al*., 2025). Improving quantitative link prediction therefore remains an important methodological frontier. Future efforts could explore alternative strategies for aggregating and transferring link weights across space, for instance by integrating abundance-based scaling or probabilistic weighting schemes, and by employing other modelling approaches than matrix factorisation.

Finally, the shared latent space assumption may be limiting in systems spanning strong environmental or phylogenetic gradients, where trait filtering and phylogenetic conservatism impose hard constraints on which species can interact. In such systems, ecologically distant locations may not share a meaningful common latent structure, and forcing them into one could obscure rather than transfer relevant interaction signals. This limitation is empirically shown by the effect of distance decay on *F*_0.5_ we found. Nevertheless, defining the boundaries of a relevant ecological context is a common working assumption in ecology, not unique to the methods we developed. However, even within a bounded ecological system, future work could address this more formally through hierarchical or multi-level embedding approaches, in which a global SVD captures shared structural patterns while location-specific offsets account for interaction niches unique to each site.

Our framework is transferable and scalable to a wide range of systems with spatial or temporal replication, requiring only moderate computation because soft-thresholded SVD operates efficiently on sparse matrices. To illustrate this generality, we applied the pipeline to both a spatially replicated island pollination system and a temporally replicated host–parasite network, demonstrating that the framework performs consistently across different ecological contexts and types of replication. Future extensions should consider three elements: (i) the spatial or temporal scale of the networks, and the resulting similarity between them; (ii) strategies for matrix aggregation; and (iii) approaches to link withholding. In this study we explored several of these dimensions — including spatial scale, alternative link withholding strategies, and sensitivity to model parameters — but the full range of possible extensions remains a rich avenue for future work. We primarily tested our approach on an island system with clear spatial separation between locations, where differences among islands are expected due to geological history (Trøjelsgaard *et al*., 2013, 2015). Natural next steps include extending the approach to inland mosaics, such as natural habitat patches within agricultural landscapes, to evaluate how spatial signals vary with habitat connectivity. Less evident extensions are applications to directional gradients such as elevation and climate, where unlike space, source–target predictions would need to be constrained to move along the gradient direction. Furthermore, our dataset comprises a relatively small number of networks with moderate species richness; whether the distance-decay signal and the benefits of auxiliary information generalise to larger, more species-rich systems remains an open question.

In conclusion, our study shows that latent patterns in network structure can effectively capture spatial signals, enabling the evaluation of how external data can improve the prediction of missing ecological interactions. Our results demonstrate that our ability to predict these missing links is spatially variable. Therefore, while network structure is highly informative for prediction, incorporating spatial context is essential to determine when and where this information can be reliably applied. The flexible framework we introduce offers an opportunity to integrate spatial variability into link prediction, supporting more realistic and ecologically informed predictions across diverse replicable systems.

## 5 Code and data availability

The data are available in the repository set up in the original publications by Trøjelsgaard *et al*. (2015): https://datadryad.org/dataset/doi:10.5061/dryad.76173 and by Pilosof *et al*. (2013): https://datadryad.org/dataset/doi:10.5061/dryad.d3d36. The full code and technical descriptions on how to run our pipeline are available on the GitHub repository https://github.com/Ecological-Complexity-Lab/svd_based_spatial_prediction/tree/clean_for_publish.

## 6 Acknowledgments

We are thankful to P. Jordano and A. Valido for their valuable feedback.

## 8 Author contribution statements

**Kesem Abramov** conceptualized the study, developed the methodology, conducted the investigation, performed formal analysis, developed and implemented software, prepared visualizations, wrote the original draft and revised the manuscript; **Geut Galai** developed and implemented software, performed formal analysis, prepared visualizations and conducted validation; **Barry Biton** developed the software, performed formal analysis and provided validation; **Rami Puzis** developed the methodology and reviewed manuscript; **Shai Pilosof** conceptualized the study, secured funding and resources, conducted investigation, developed the methodology, developed software, provided supervision, wrote the original draft and revised the manuscript. All authors contributed critically to the drafts and gave final approval for publication.

## 9 Supplementary information

## 9.1 Appendix S1 Scale analysis for primary case study (pollination network)

**Fig. S1:**
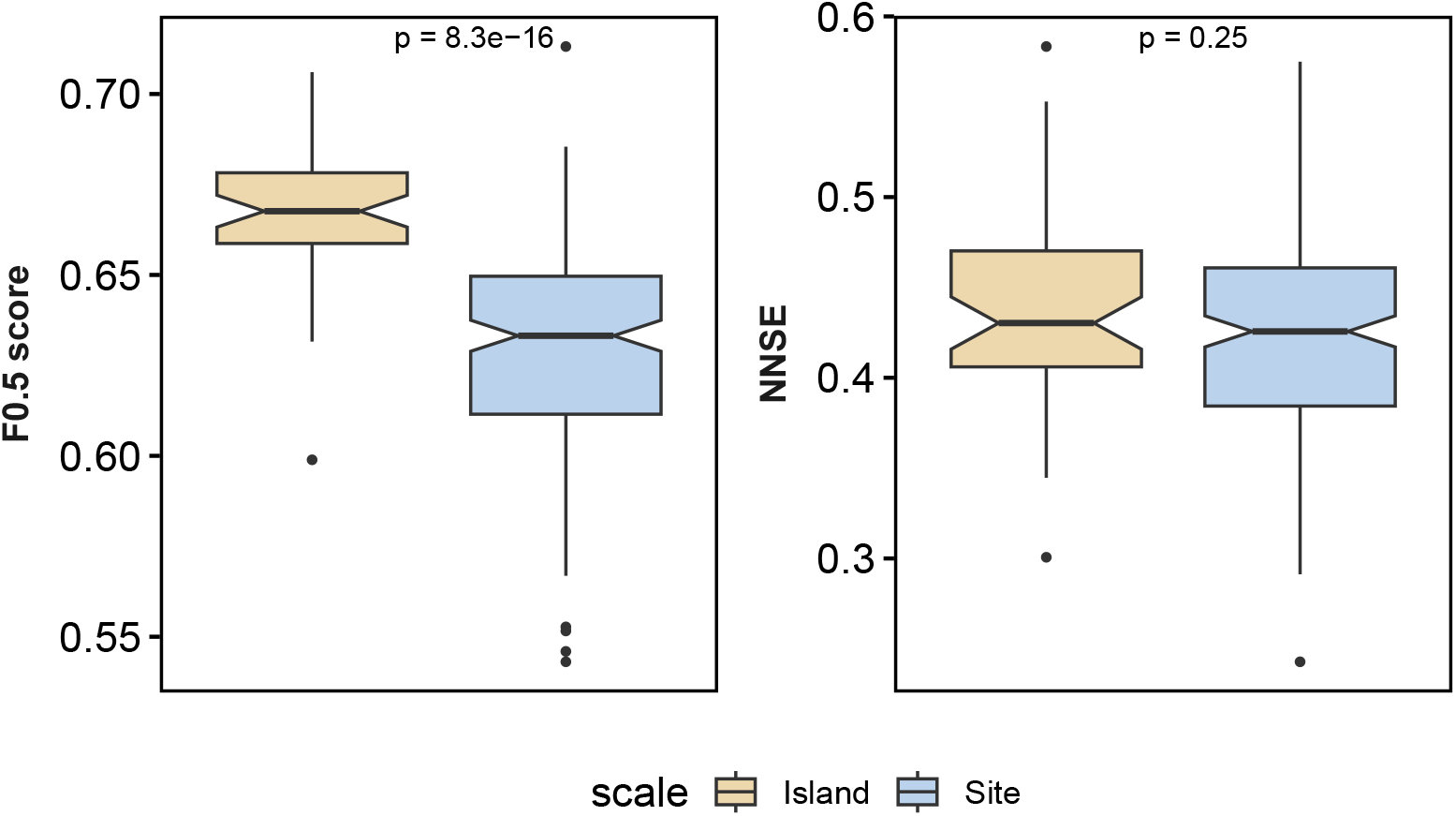
Effects of spatial scale on predictive performance. Binary link prediction capacity, measured by *F*_0.5_ scores, is higher when site scale data are aggregated into island scale. However, weighted predictive power (NNSE) is similar at the site and island scales. P values denote Wilcoxon test results.

**Fig. S2:**
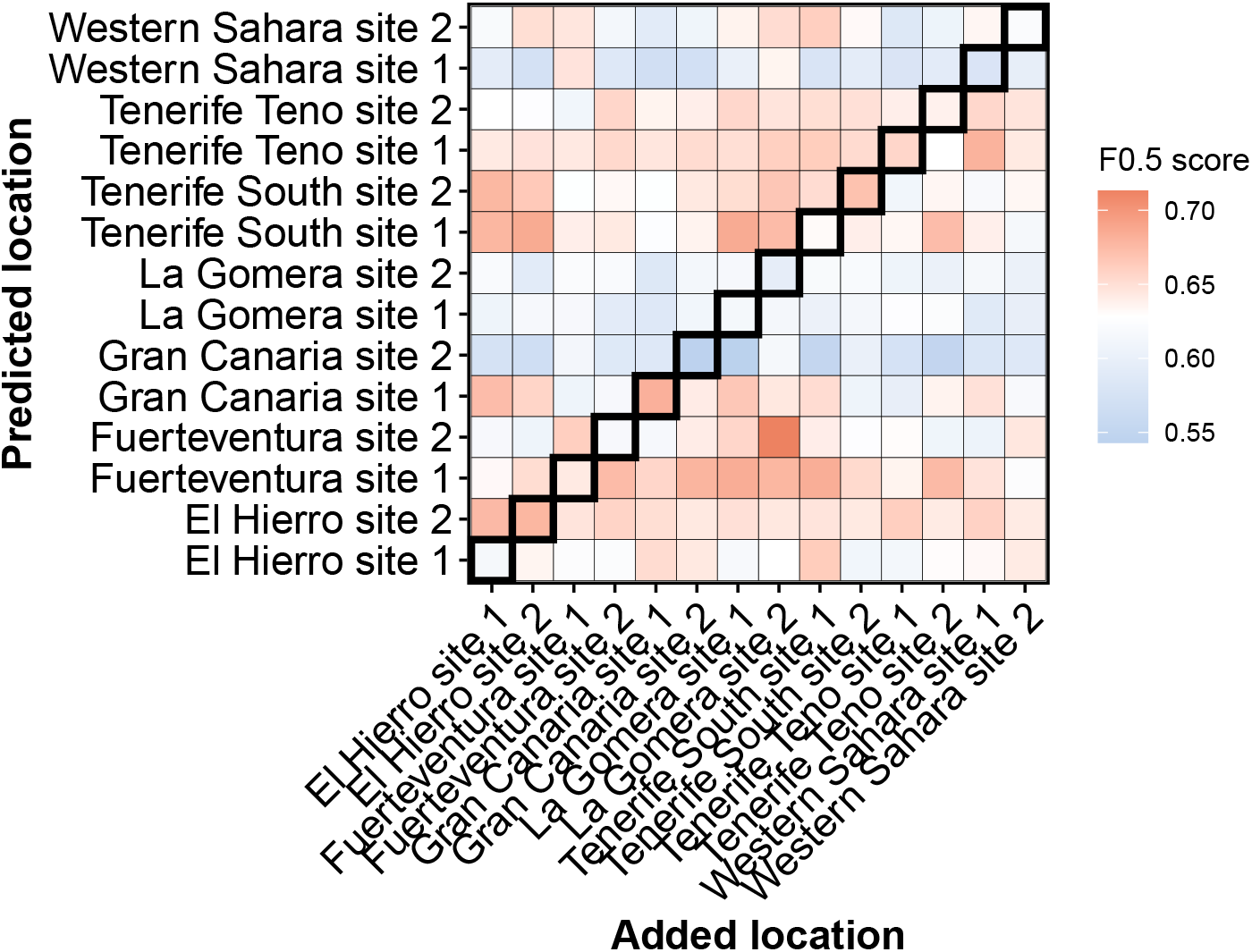
Site-scale variation in predictive capacity. *F*_0.5_ scores vary according to the pair of predicted and added sites.

**Fig. S3:**
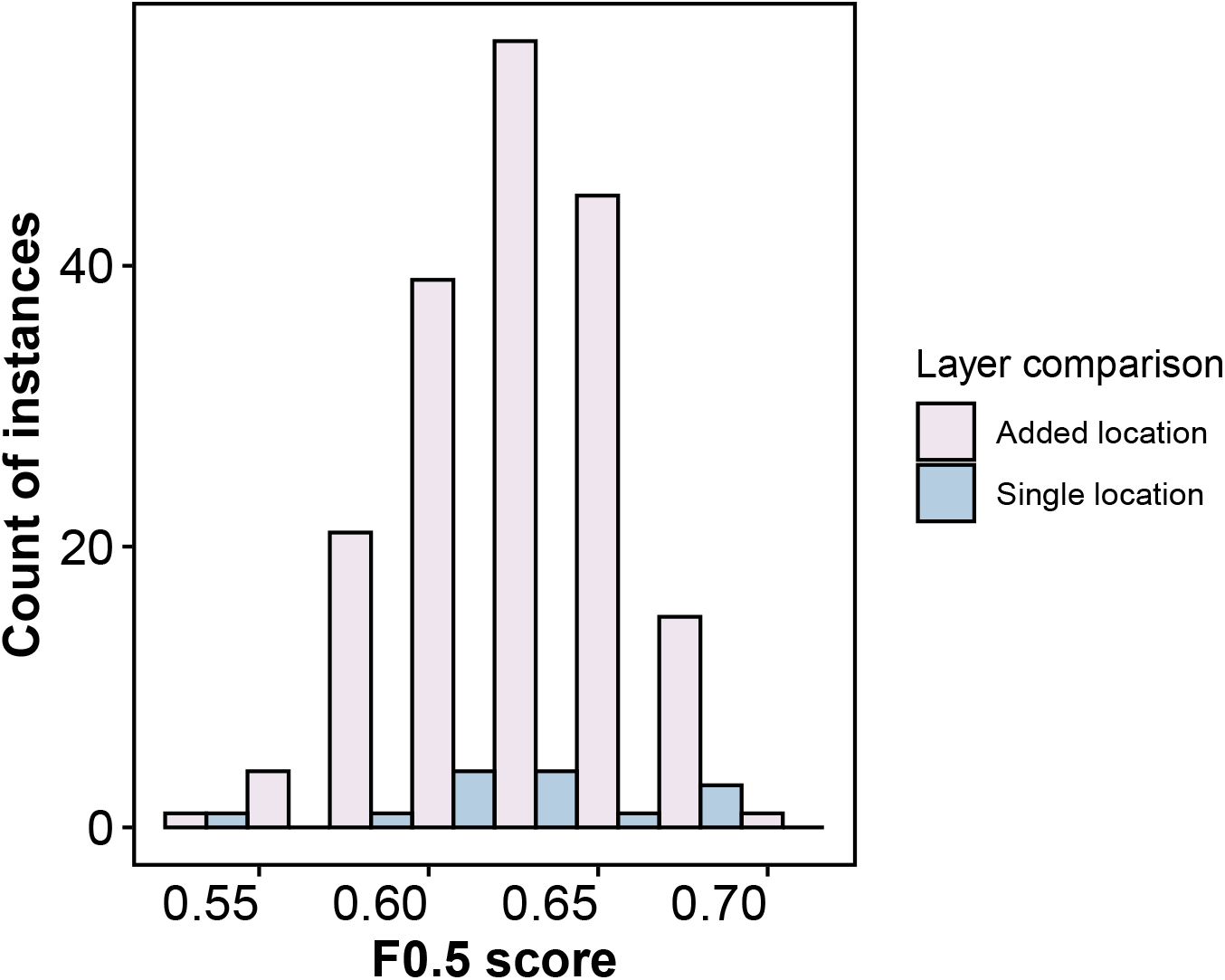
Distribution of site-scale prediction quality. Quality of binary predictions according to the *F*_0.5_ scores when incorporating data from other sites compared to using single-site data.

**Fig. S4:**
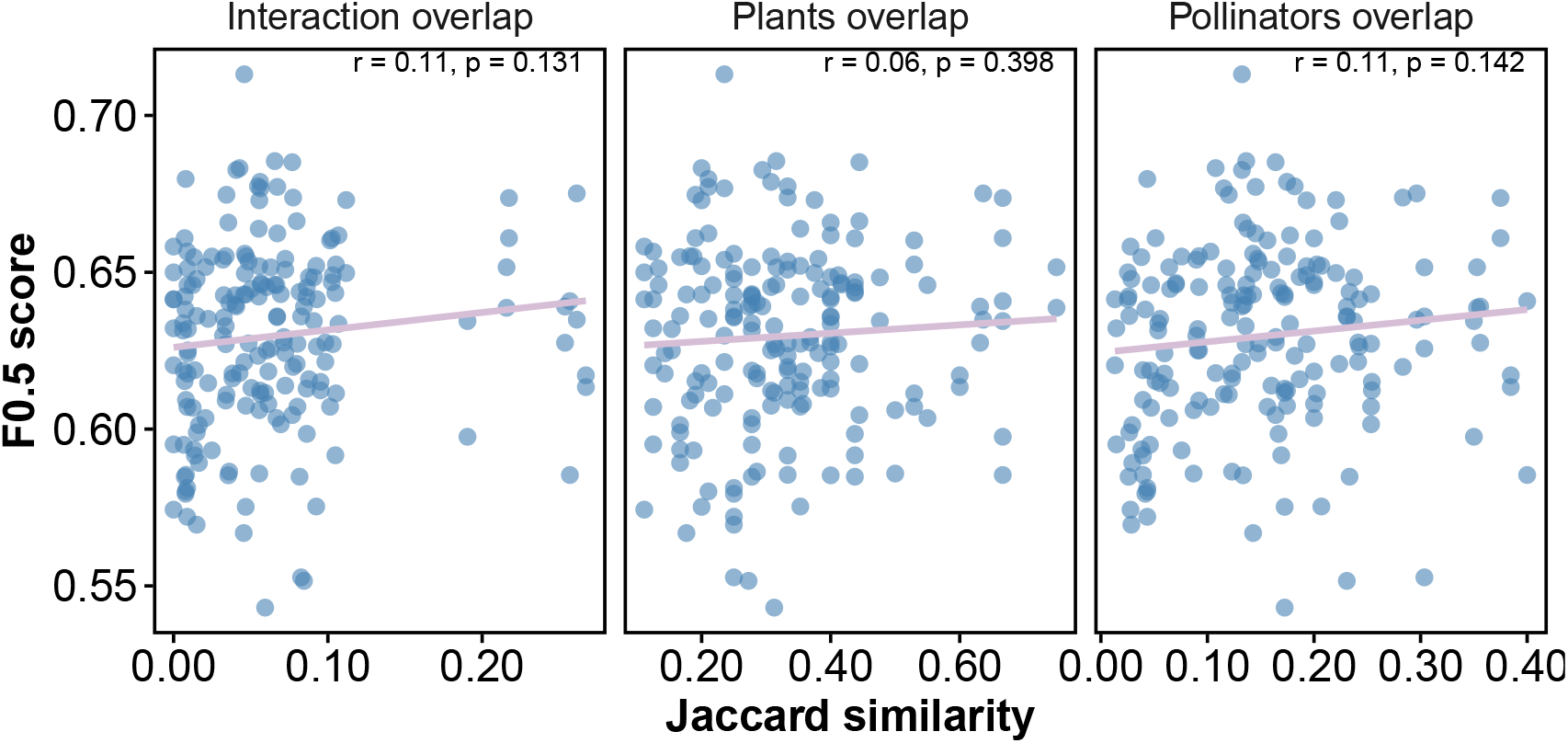
Site-scale predictive quality and ecological similarity between sites. While there are weak positive trends, the relationship between predictive ability (*F*_0.5_ scores) and ecological similarity (in interaction and species composition) between predicted and added sites is not significant.

**Fig. S5:**
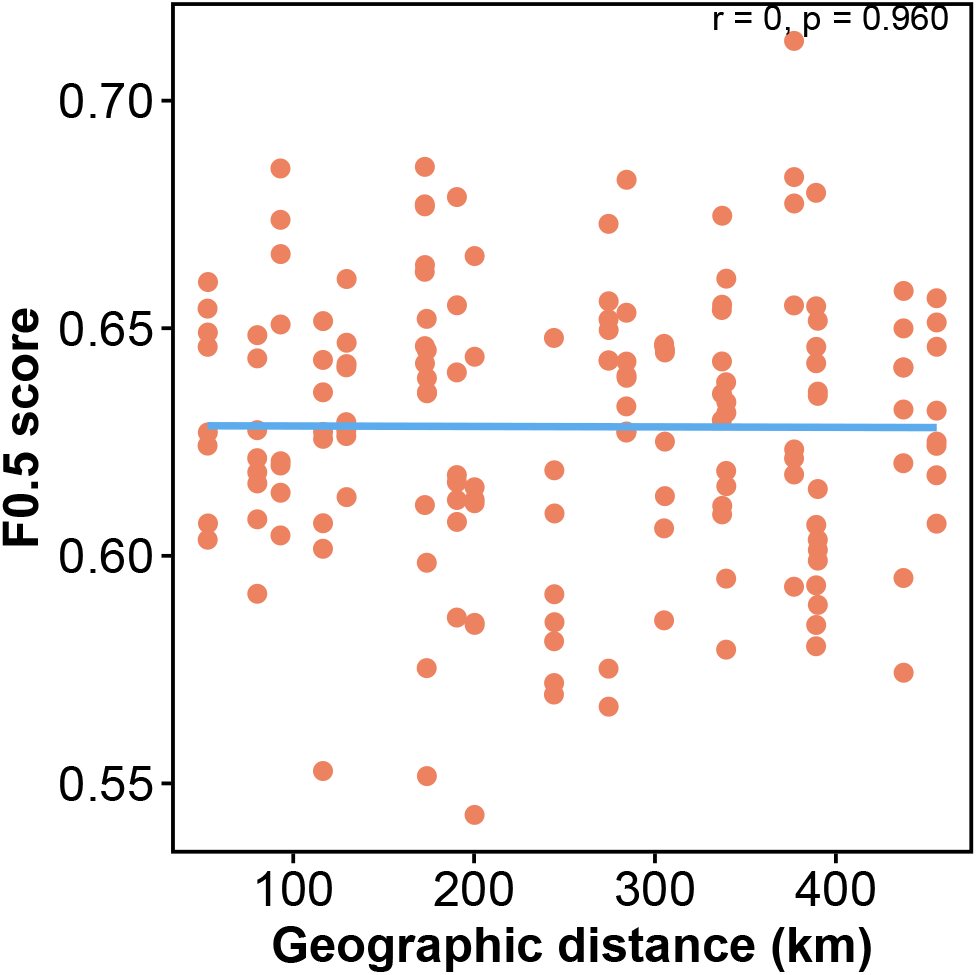
Distance decay not detected in site-scale predictive performance. Predictive accuracy does not show a correlation with geographical distance between predicted and added sites, aligned with MRM test results (*P* = 0.56).

**Fig. S6:**
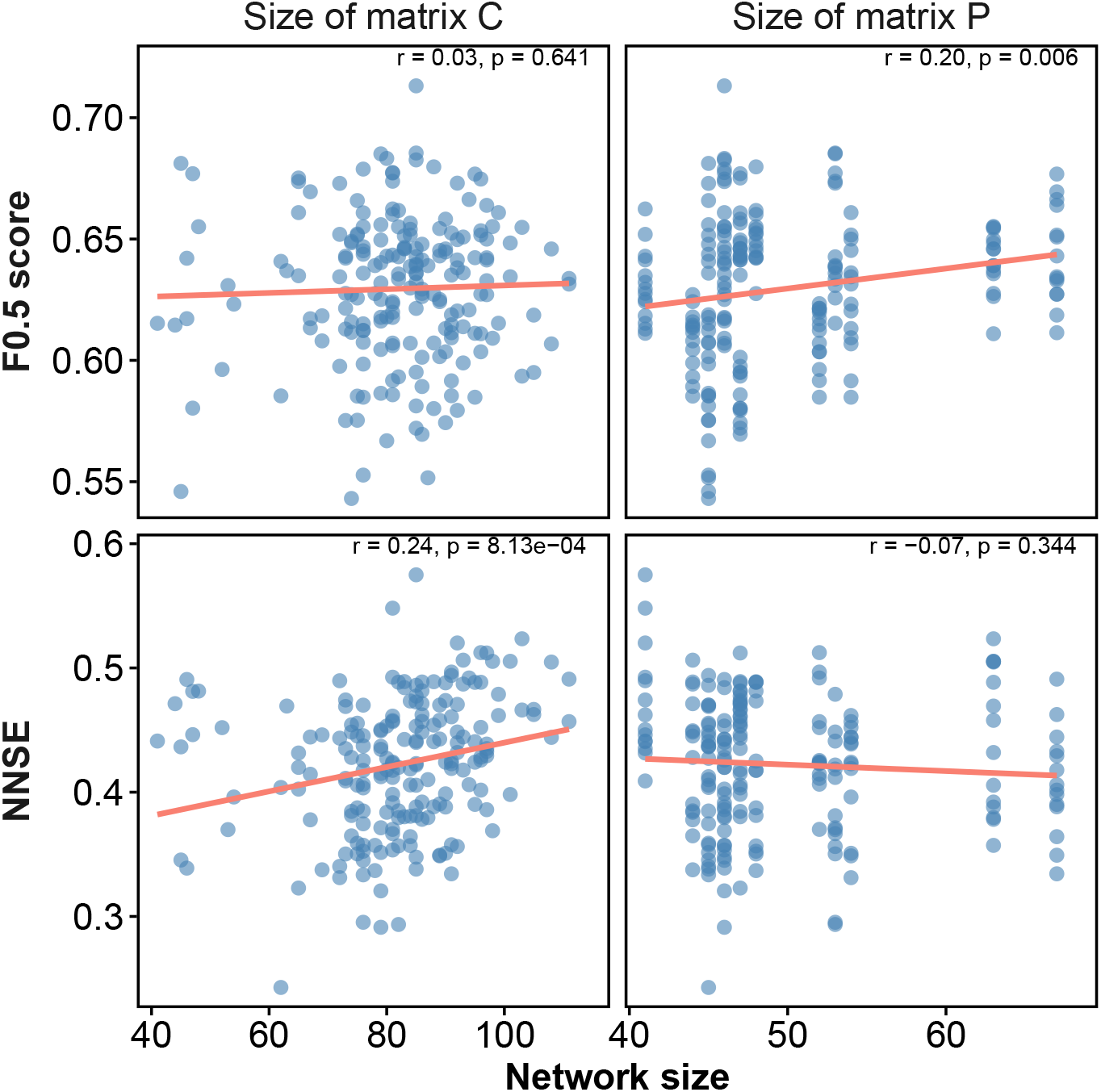
Effect of site-scale network size on predictive ability. Binary prediction quality (upper panels) tends to improve as the size of the predicted network increases. Weighted prediction quality, on the other hand, increases as the size of the matrix combining interactions from the predicted and added sites increases (lower panels). Nevertheless, the effect sizes are small.

**Fig. S7:**
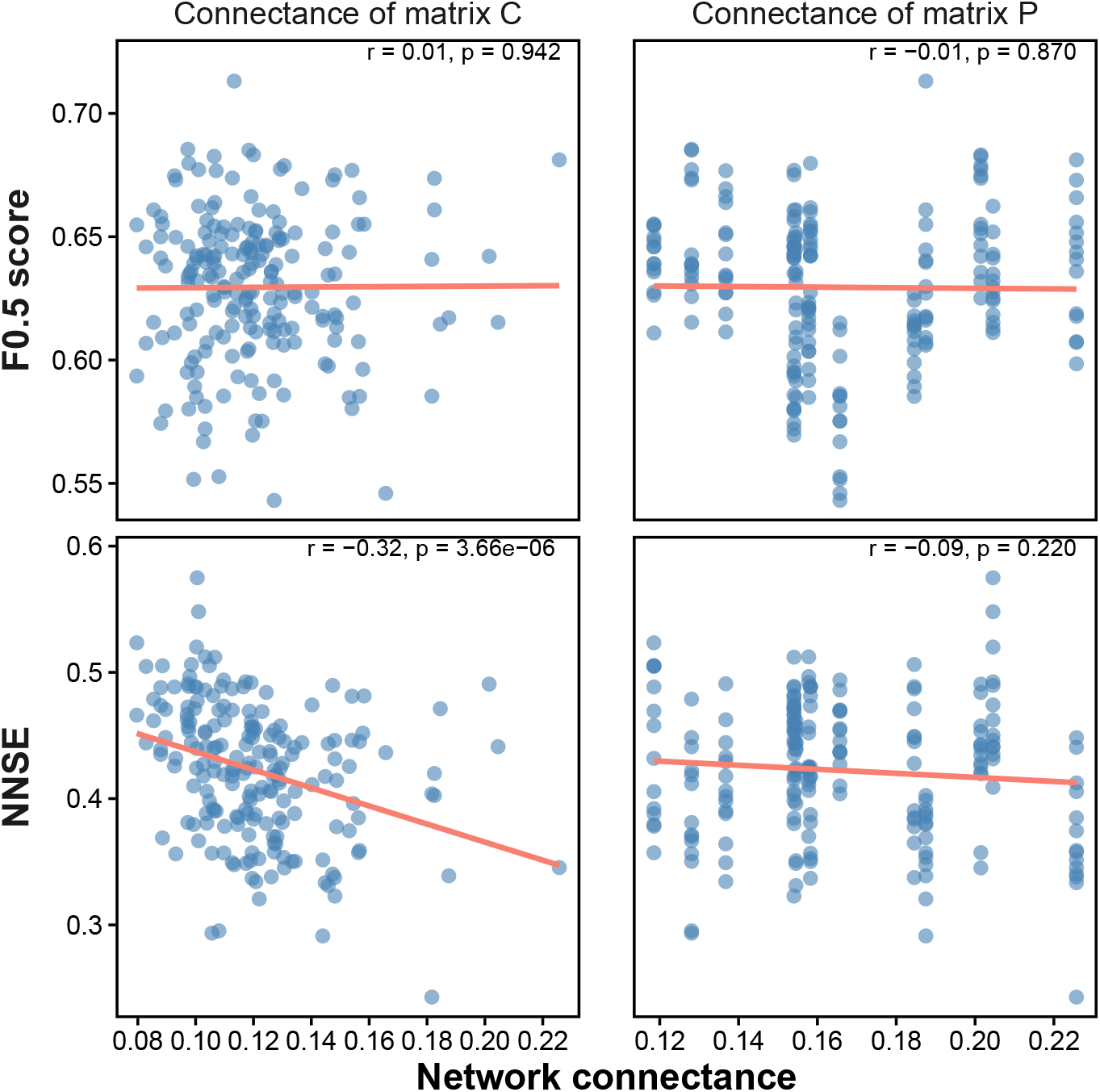
Effect of site-scale network connectance on predictive performance. Weighted (but not binary) prediction performance tends to decrease with increasing density of the combined matrix (left panels), whereas both binary and weighted prediction quality show no relationship with the density of the target matrix (right panels).

## 9.2 Appendix S2 Additional analyses of the main case study (pollination network)

**Fig. S8:**
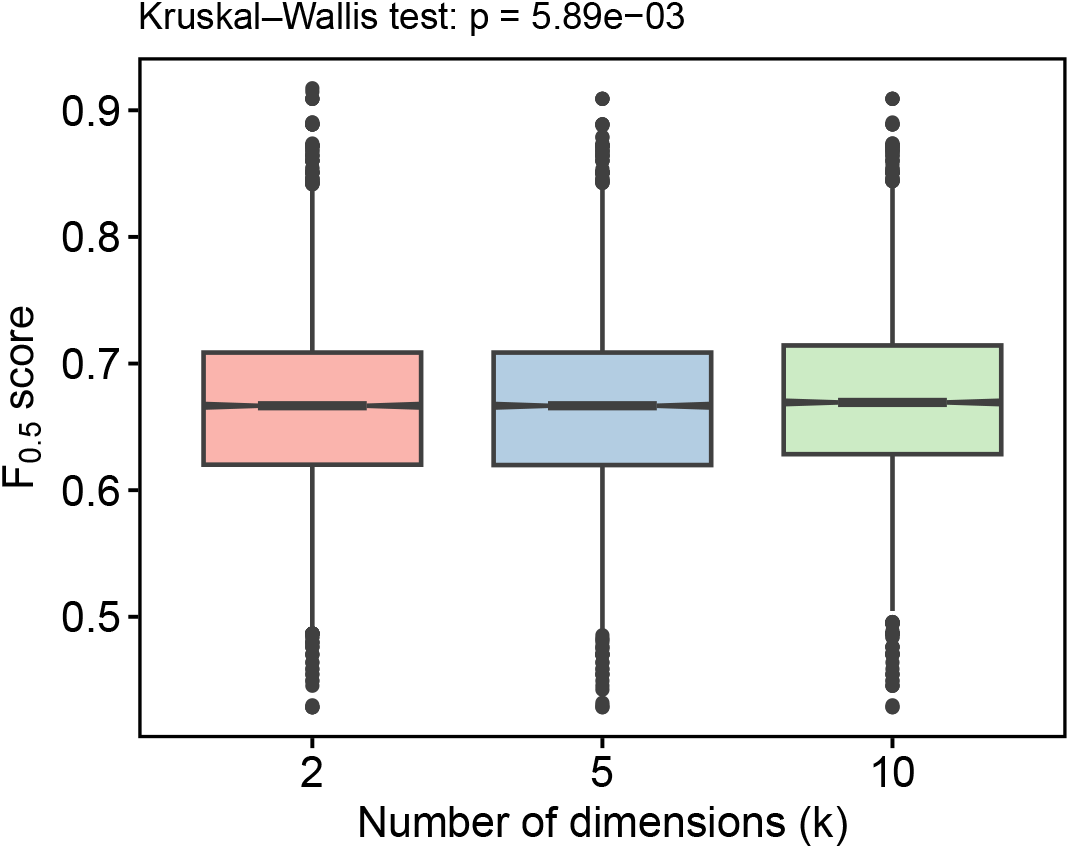
Sensitivity analysis for the maximal number of latent dimensions (*k*). Prediction performance (F_0.5_) was largely insensitive to the choice of *k*. Although differences among *k* values were statistically significant due to the large sample size (*n* = 29400 model evaluations across all *k* values, *λ* values, networks and withholding iterations), the effect sizes were negligible (Cohen’s *d <* 0.04, i.e., minimal differences in means relative to the pooled standard deviation across all pairwise comparisons of *k* values), indicating that the main structural signal is captured with few latent dimensions.

**Fig. S9:**
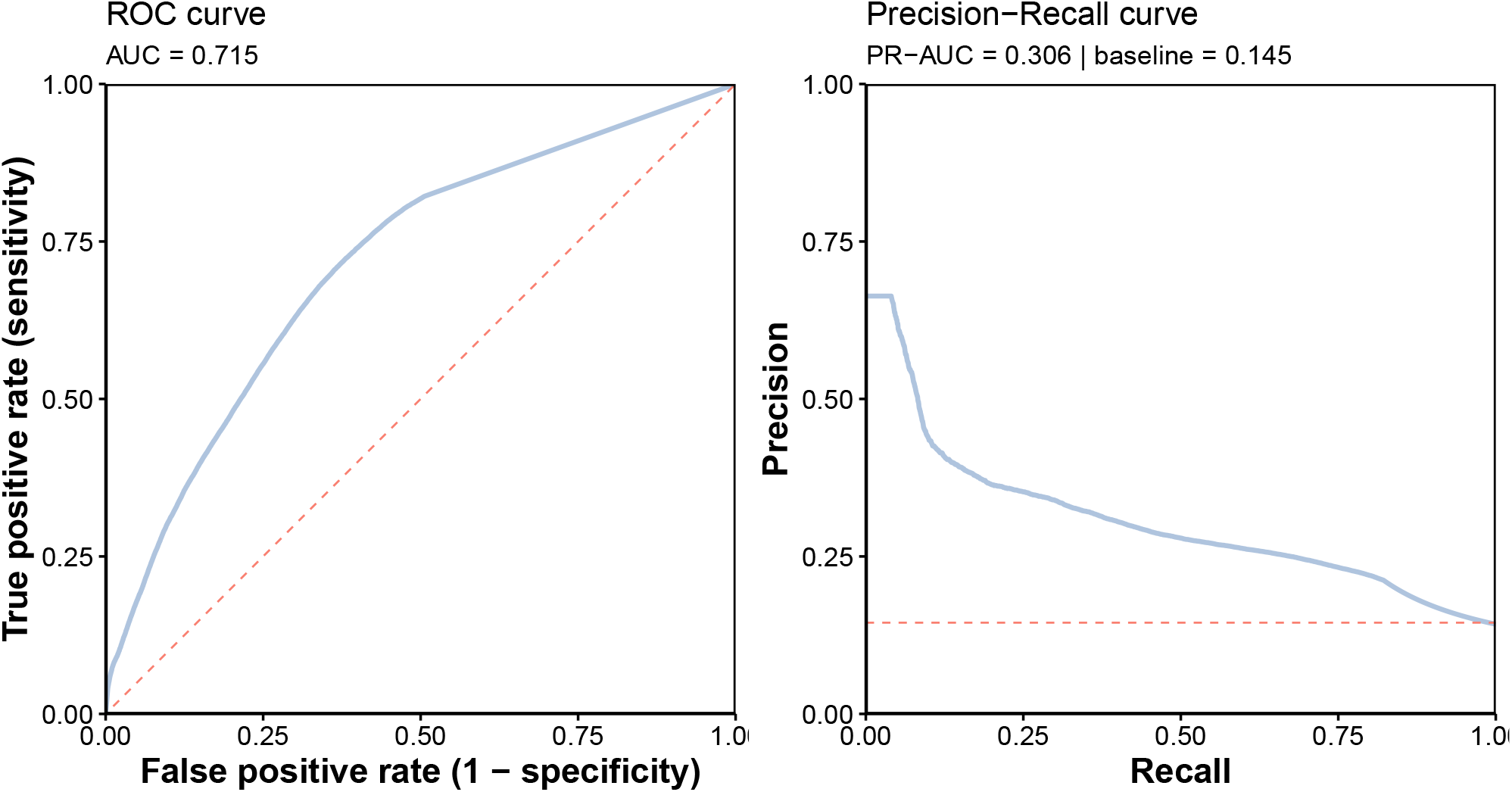
Predictive performance under class-imbalance-preserving link withholding. Receiver operating characteristic (ROC) and precision–recall (PR) curves obtained when links and non-links are withheld according to their prevalence in the interaction matrices (across all prediction cases and classification thresholds). The dashed diagonal line in the ROC panel indicates random performance. In the PR panel, the dashed horizontal line shows the baseline precision equal to the prevalence of true links in the evaluation set. These curves show that when links and non-links are withheld proportionally to their prevalence, predictive performance remains well above the random baseline, indicating that the algorithm performs well even under the class imbalance typical of ecological interaction networks.

**Fig. S10:**
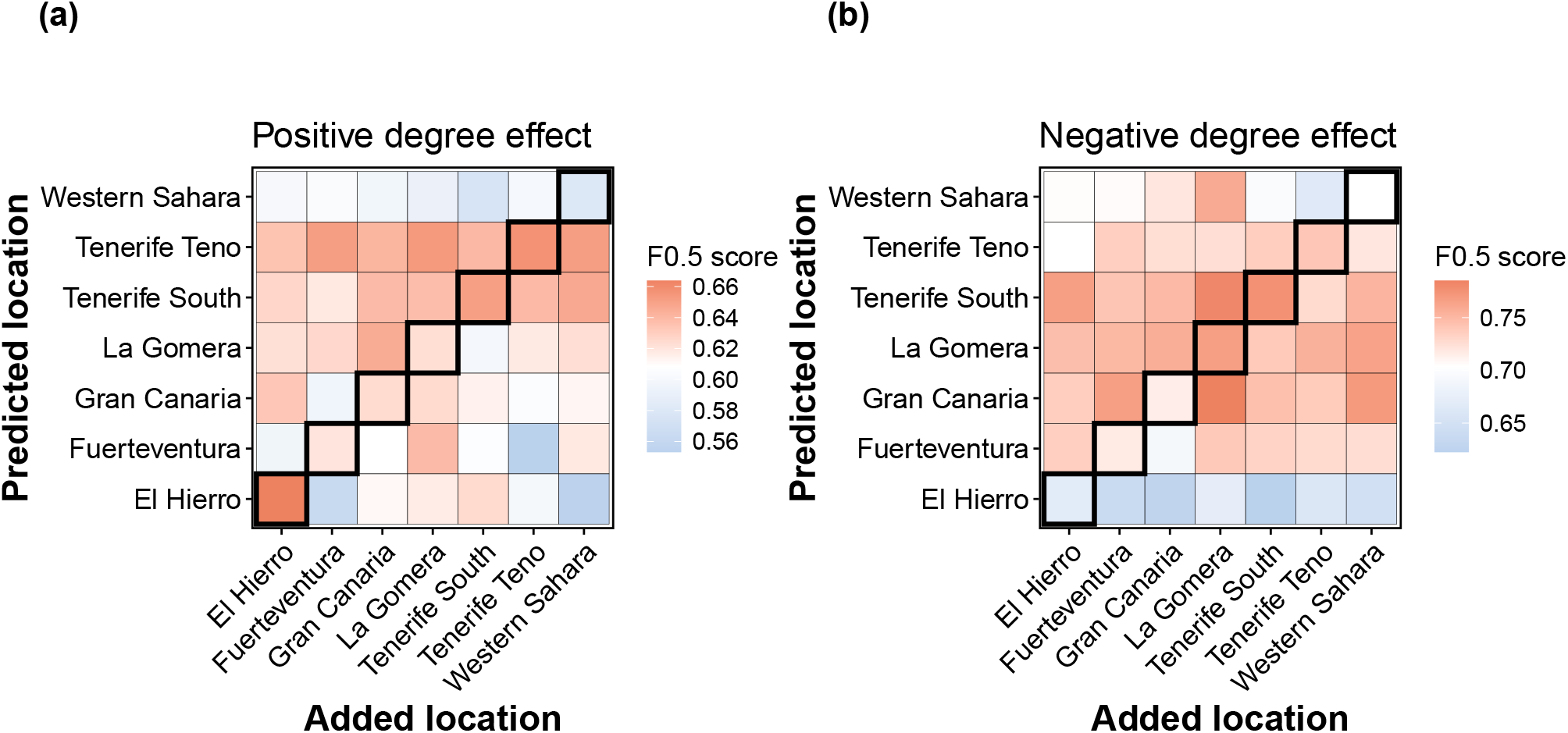
Influence of link withholding strategy on predictive performance. Predictive performance across 49 prediction cases using each of the seven locations as target or auxiliary networks when links are withheld **(a)** with positive association to the degrees of the species involved (i.e., interactions involving generalists are preferentially withheld) and **(b)** with negative association to species’ degrees (i.e., interactions involving specialists are preferentially withheld).

**Table S1:** Performance metrics used to evaluate model predictions. TP, FP, FN and TN denote true positives, false positives, false negatives, and true negatives, respectively. *y*_*i*_ and *ŷ*_*i*_ denote observed and predicted values, and 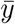 is the mean of observed values.

| Metric | Formula | Definition |
| --- | --- | --- |
| **Precision** | $\frac{TP}{TP + FP}$ | Proportion of predicted interactions that are true. |
| **Recall** | $\frac{TP}{TP + FN}$ | Proportion of true interactions correctly predicted. |
| **$F_{0.5}$ score** | $1.25 \cdot \frac{\text{Precision} \cdot \text{Recall}}{0.25 \cdot \text{Precision} + \text{Recall}}$ | Weighted harmonic mean of precision and recall, giving more weight to precision (thus penalising false positives more strongly than false negatives). |
| **$F_1$ score** | $2 \cdot \frac{\text{Precision} \cdot \text{Recall}}{\text{Precision} + \text{Recall}}$ | Harmonic mean of precision and recall. |
| **ROC–AUC** | $\int_0^1 \text{TPR} d(\text{FPR})$ | Area under the Receiver Operating Characteristic curve across classification thresholds. Here, $\text{TPR} = \frac{TP}{TP + FN}$ and $\text{FPR} = \frac{FP}{FP + TN}$ (false positive rate). |
| **PR–AUC** | $\int_0^1 \text{Precision} d(\text{Recall})$ | Area under the Precision–Recall curve across classification thresholds. |
| **NNSE** | $\text{NNSE} = \frac{1}{2 - \text{NSE}}$ $\text{NSE} = 1 - \frac{\sum_{i=1}^n (y_i - \hat{y}_i)^2}{\sum_{i=1}^n (y_i - \bar{y})^2}$ | Normalised Nash–Sutcliffe efficiency; measures weighted predictive skill relative to predicting the mean of observed values, rescaled to the interval $[0, 1]$. |

**Table S2:** Network size and connectance (density) for each location in the data set.

| Island | Size | Connectance |
| --- | --- | --- |
| Tenerife South | 84 | 0.128 |
| Tenerife Teno | 78 | 0.120 |
| Western Sahara | 72 | 0.146 |
| La Gomera | 67 | 0.149 |
| El Hierro | 65 | 0.148 |
| Fuerteventura | 65 | 0.183 |
| Gran Canaria | 62 | 0.182 |

**Fig. S11:**
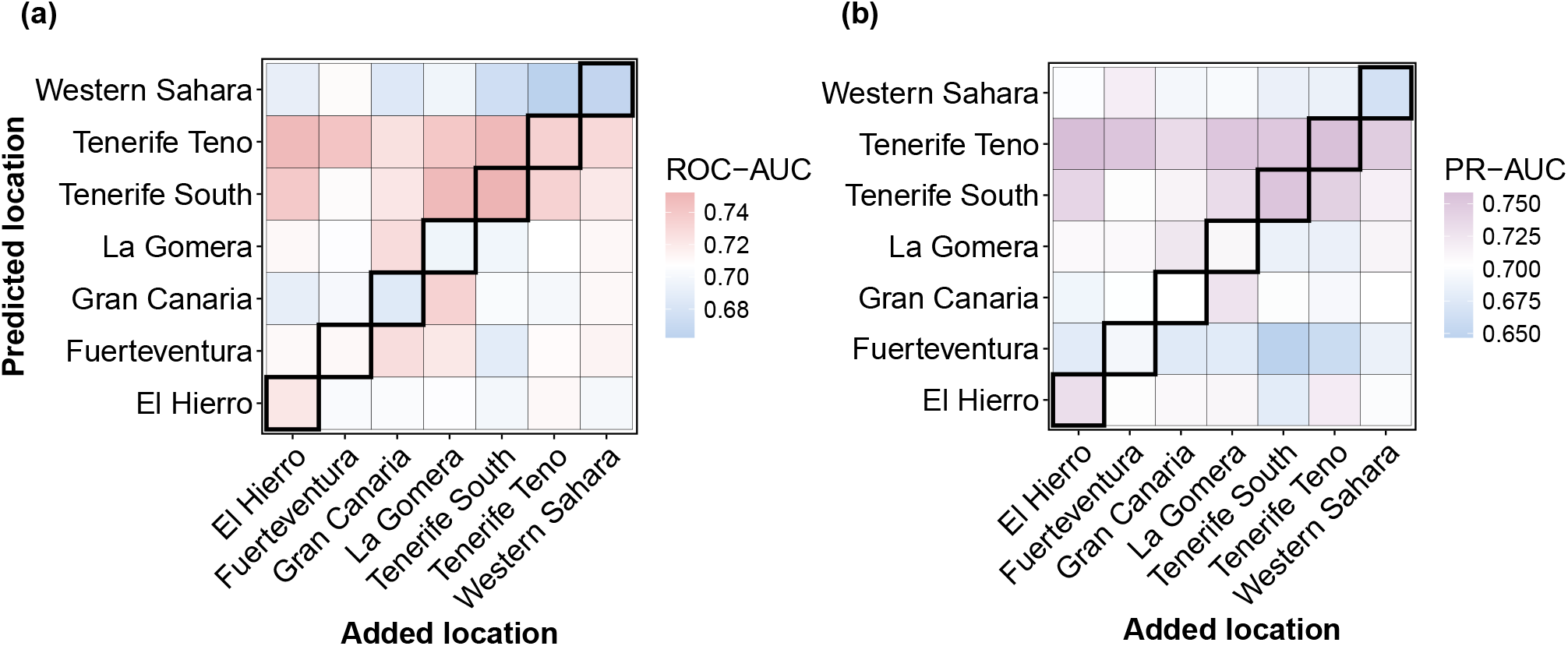
Spatial variability in non-thresholded binary predictive performance. **(a)** ROC-AUC values and **(b)** PR-AUC values vary according to the combination of predicted and added locations.

**Fig. S12:**
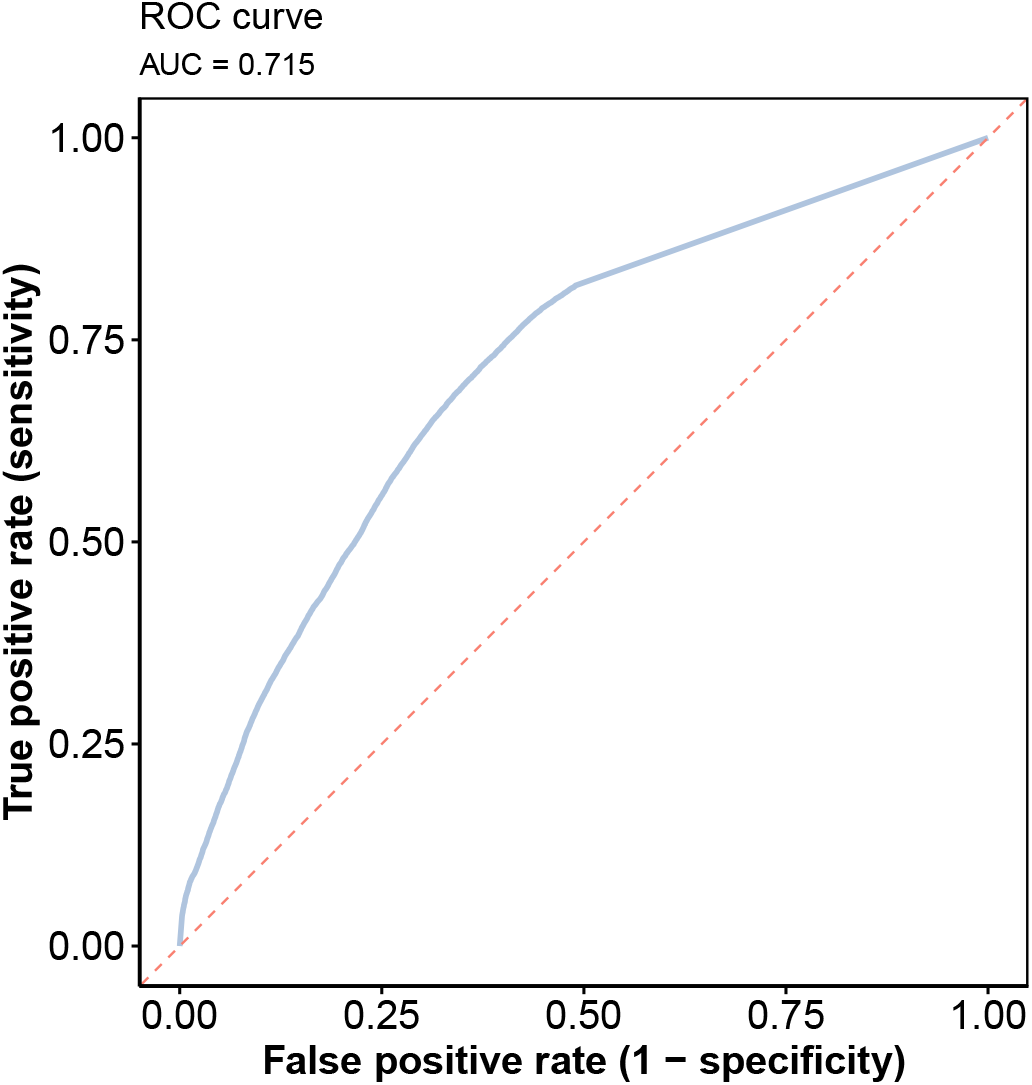
Receiver operating characteristic (ROC) curve for predicted interactions. The ROC curve shows the trade-off between the true positive rate and the false positive rate across all classification thresholds across evaluation cases (under the random withholding strategy). The dashed diagonal line represents the performance of a random predictor. The area under the curve (AUC) summarizes the algorithm’s ability to distinguish true links from non-links.

**Fig. S13:**
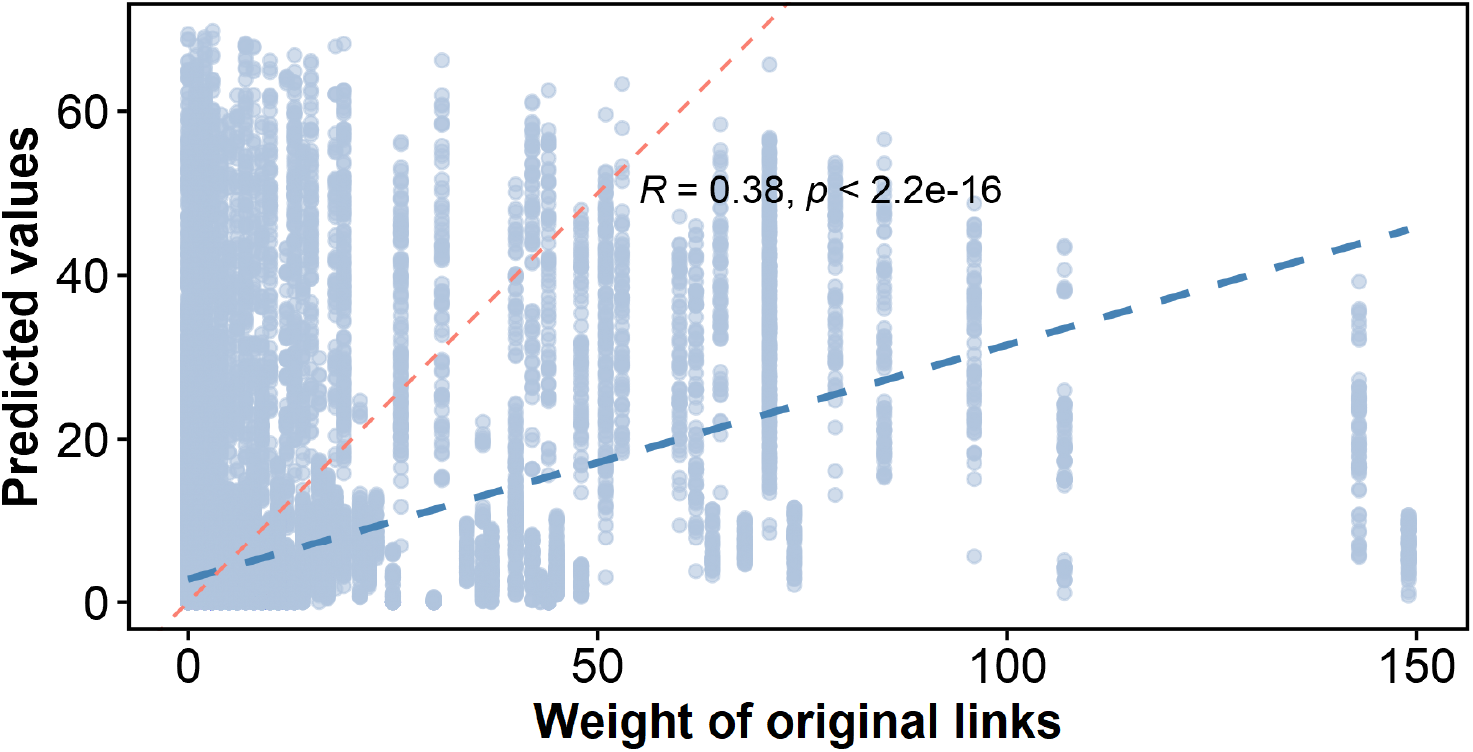
Range of predictions for observed link weights. Pearson’s correlation between predicted values and original link weights. Empiric trend line is in blue. For comparison, a trend line denoting perfect correlation is in red.

**Fig. S14:**
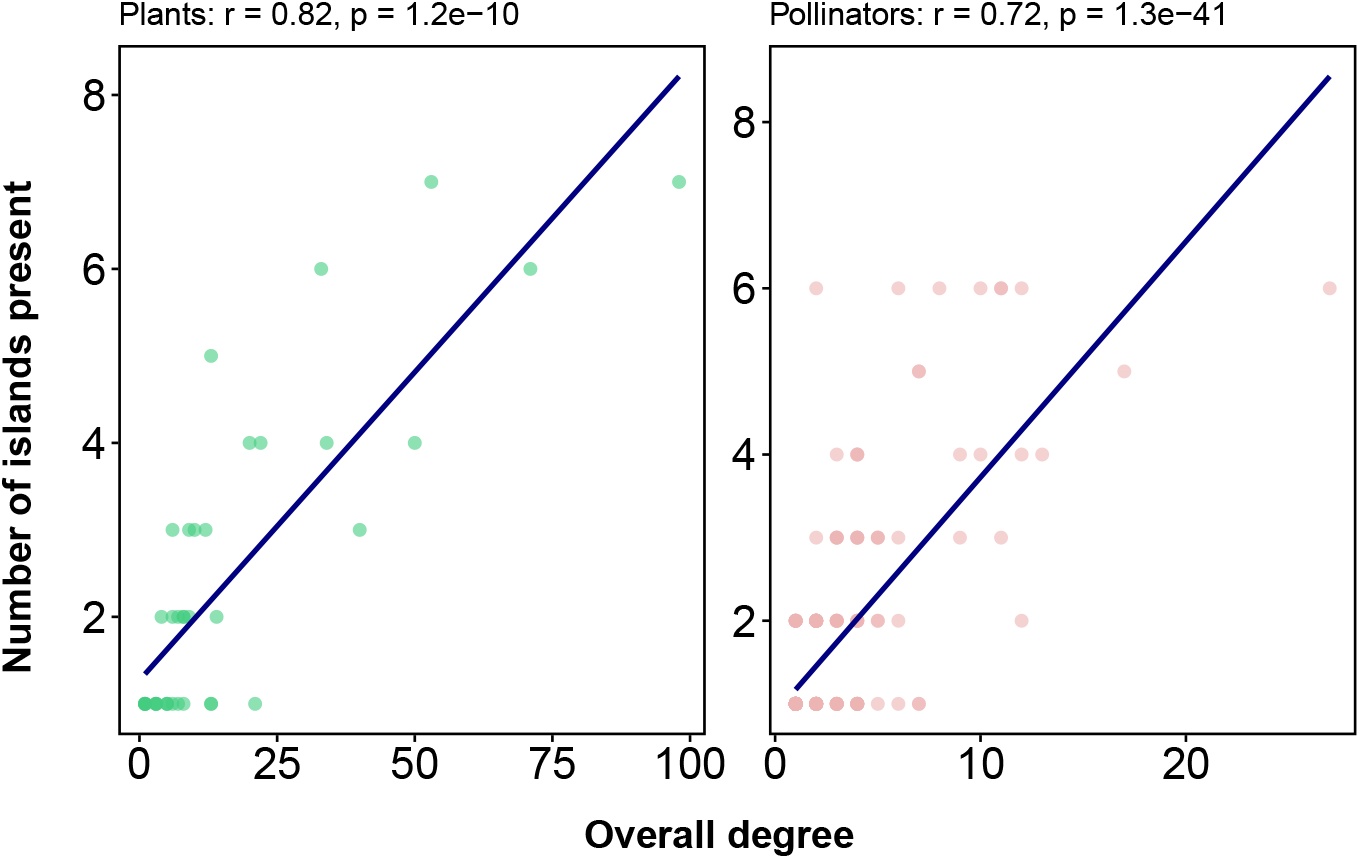
Correlation between species degree and occurrence. For each plant and pollinator species, degree (the number of unique partners across the Canary Islands, representing generalisation level) is plotted against the number of islands where the species occurs. Values denote Pearson’s correlation.

**Fig. S15:**
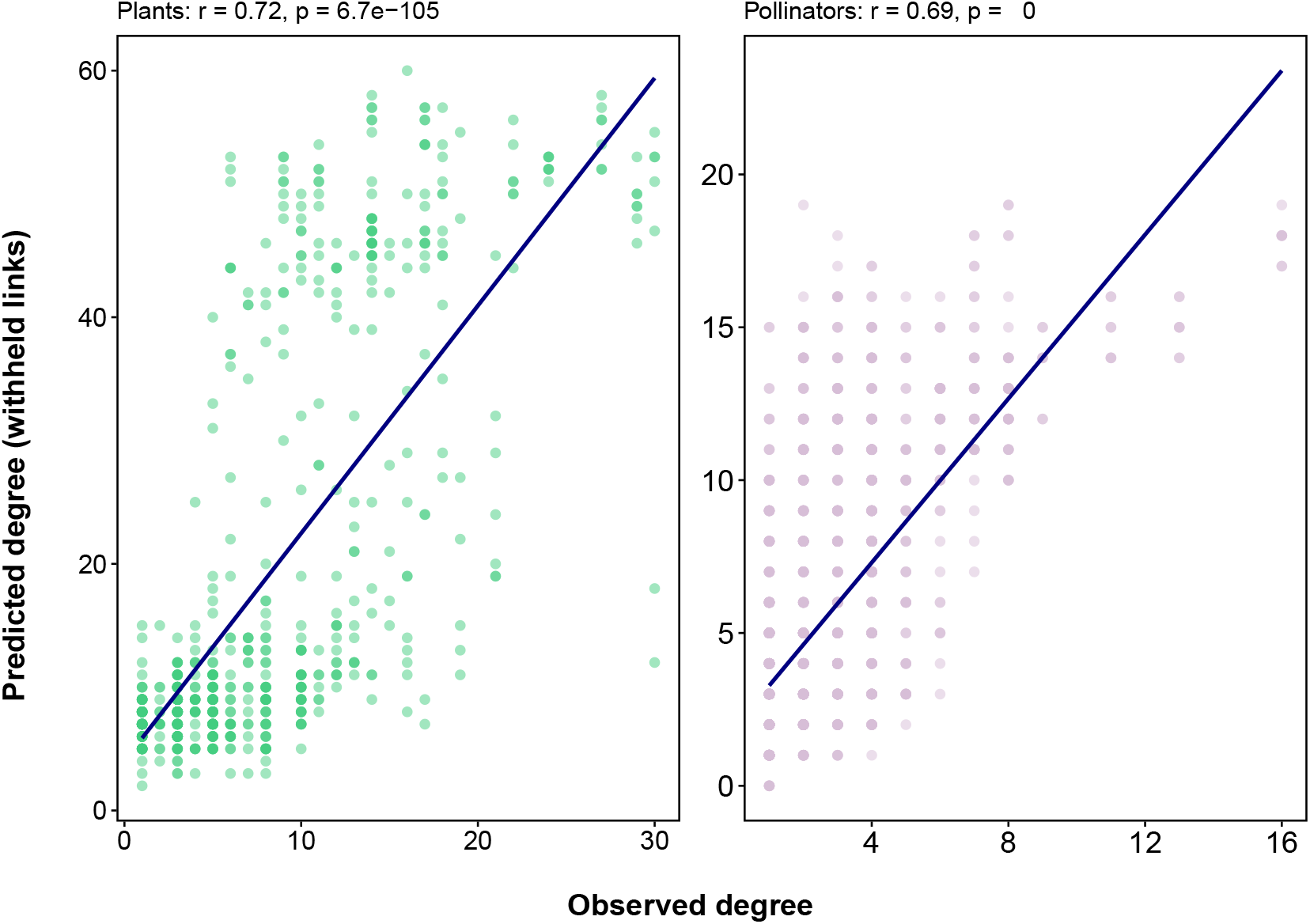
Relationship between observed and predicted local species degrees. Predicted degree corresponds to the number of predicted interactions assigned to a species in a specific combination of predicted and auxiliary networks. It is plotted against its observed degree within this specific spatial context. Although species with higher observed degree tend to receive more predicted links, the relationship is not perfect, indicating that predictions are not determined solely by degree.

**Fig. S16:**
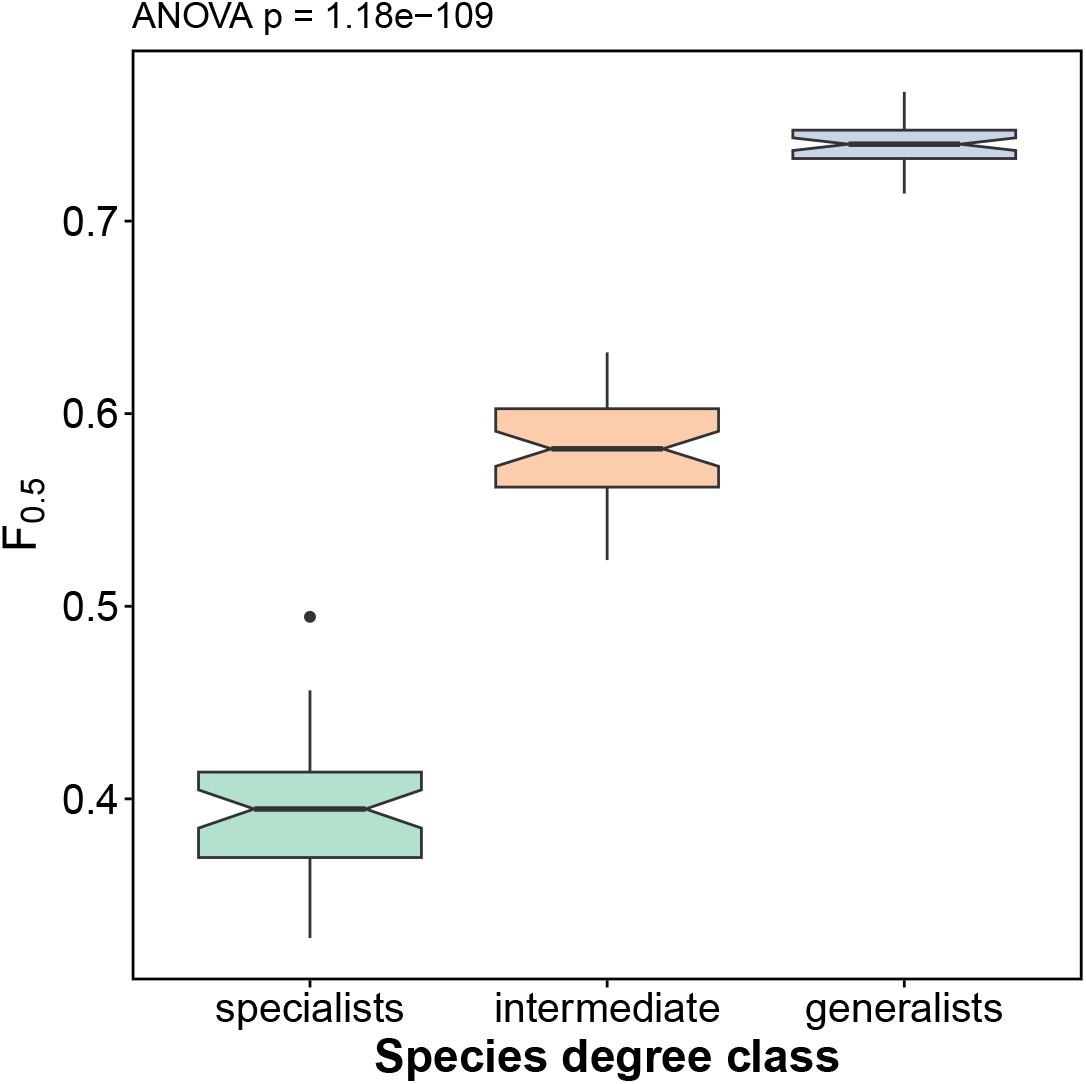
Predictive performance across species degree classes. Species were grouped according to their observed degree (third of all species, that have the lowest degrees, classified as specialists, the third with highest degrees are generalists, and the remaining third are intermediate). Boxes show the distribution of *F*_0.5_ scores across prediction cases. Predictive performance increases with species degree, reflecting the greater amount of interaction information available for high-degree species.

**Fig. S17:**
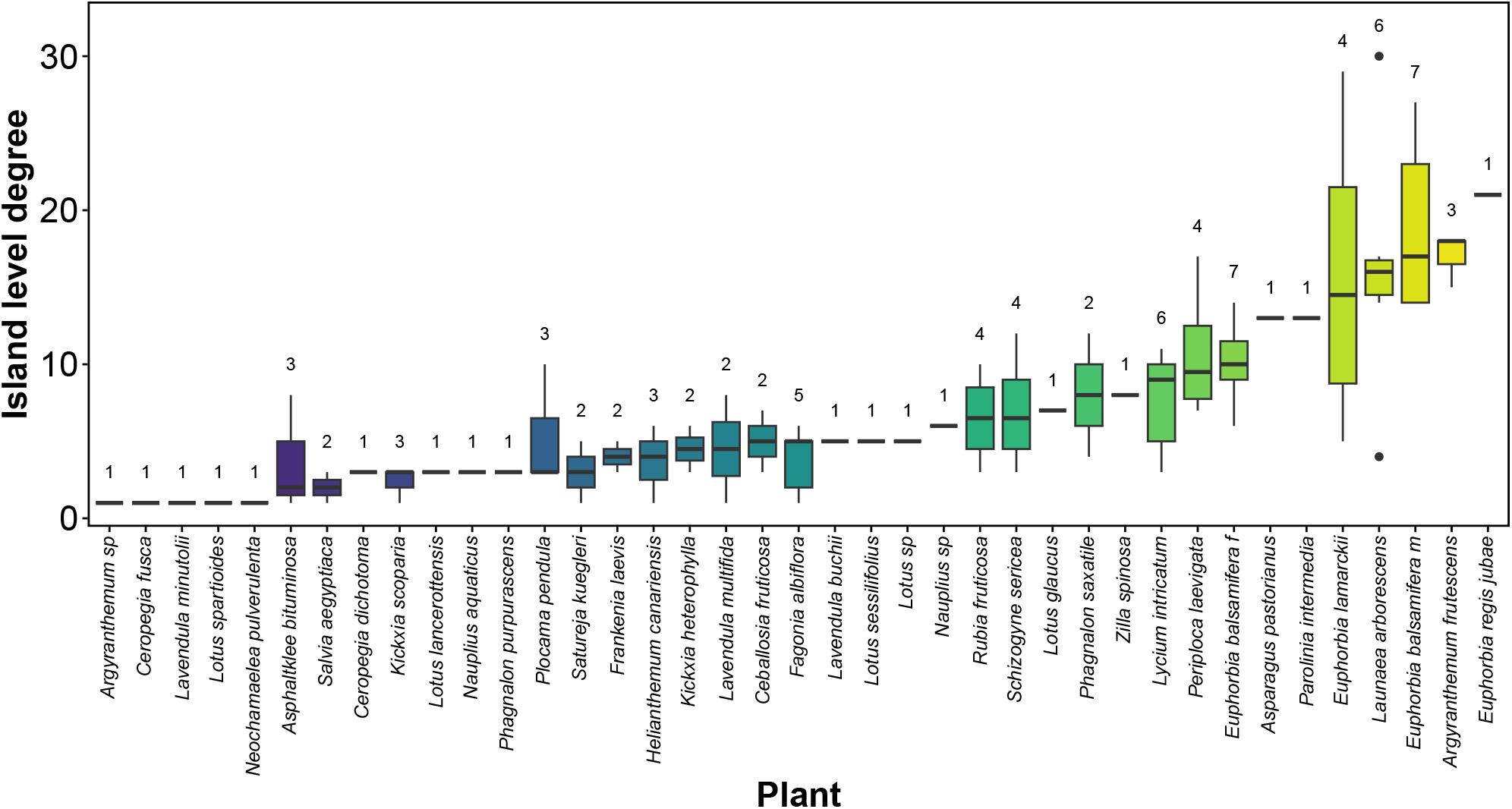
Variation in island-level degree of plant species across the system. For each plant species, boxes show the distribution of its local degree (number of interaction partners) across the islands where it occurs. The numbers above the boxes denote the number of islands in which the species occurs. With few exceptions, species that are specialists or generalists at the local level tend to show similar patterns across the islands where they occur.

**Fig. S18:**
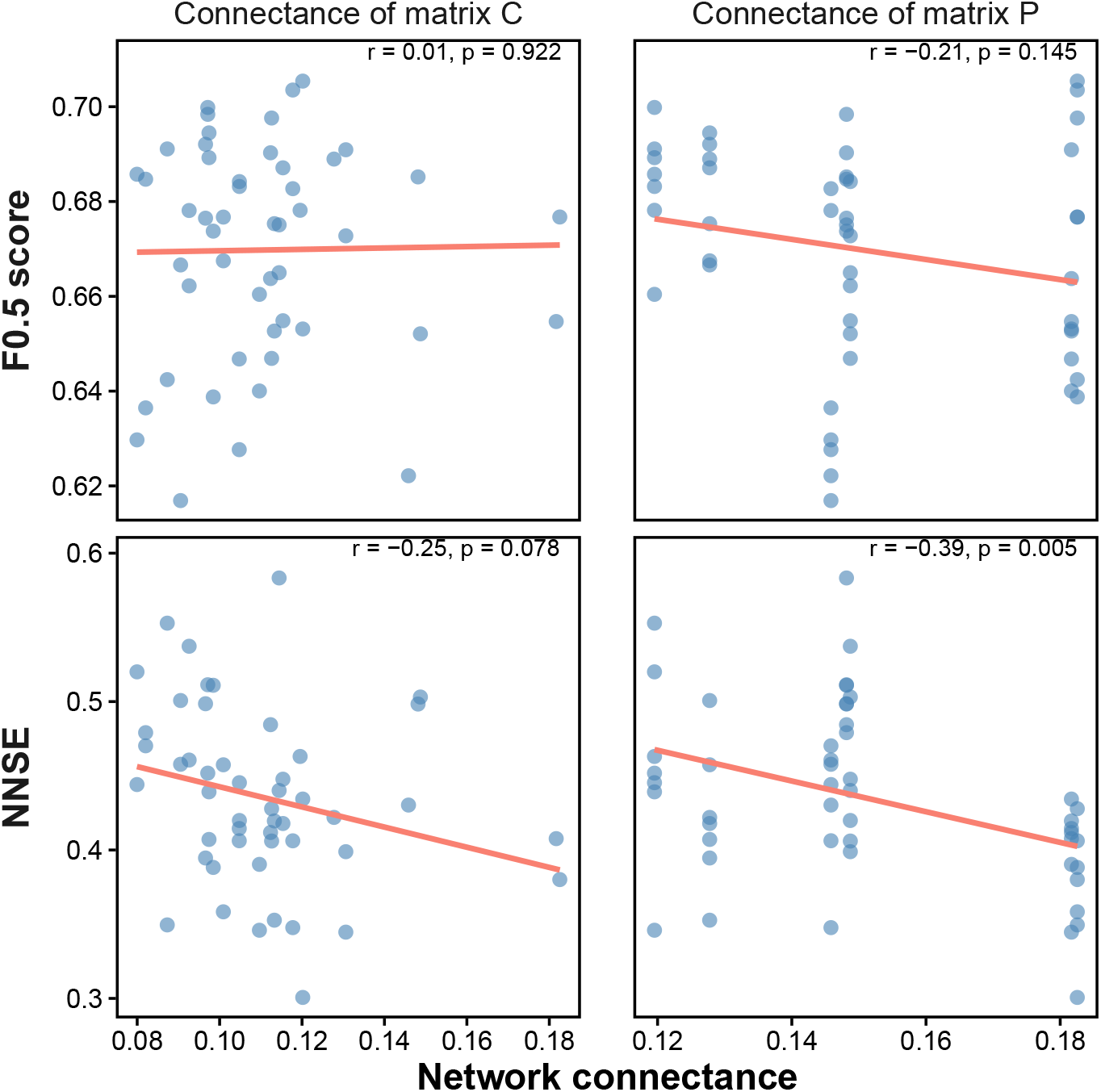
Effect of network connectance on predictive performance. Binary prediction quality was not related to the connectance of the predicted matrix **P** nor to the matrix combining interactions from the predicted and added locations (**C**) (upper panels). Weighted prediction quality significantly decreased as the target network became denser (lower right). Nevertheless, we note that for this data set weighted predictions were generally not better than random.

**Fig. S19:**
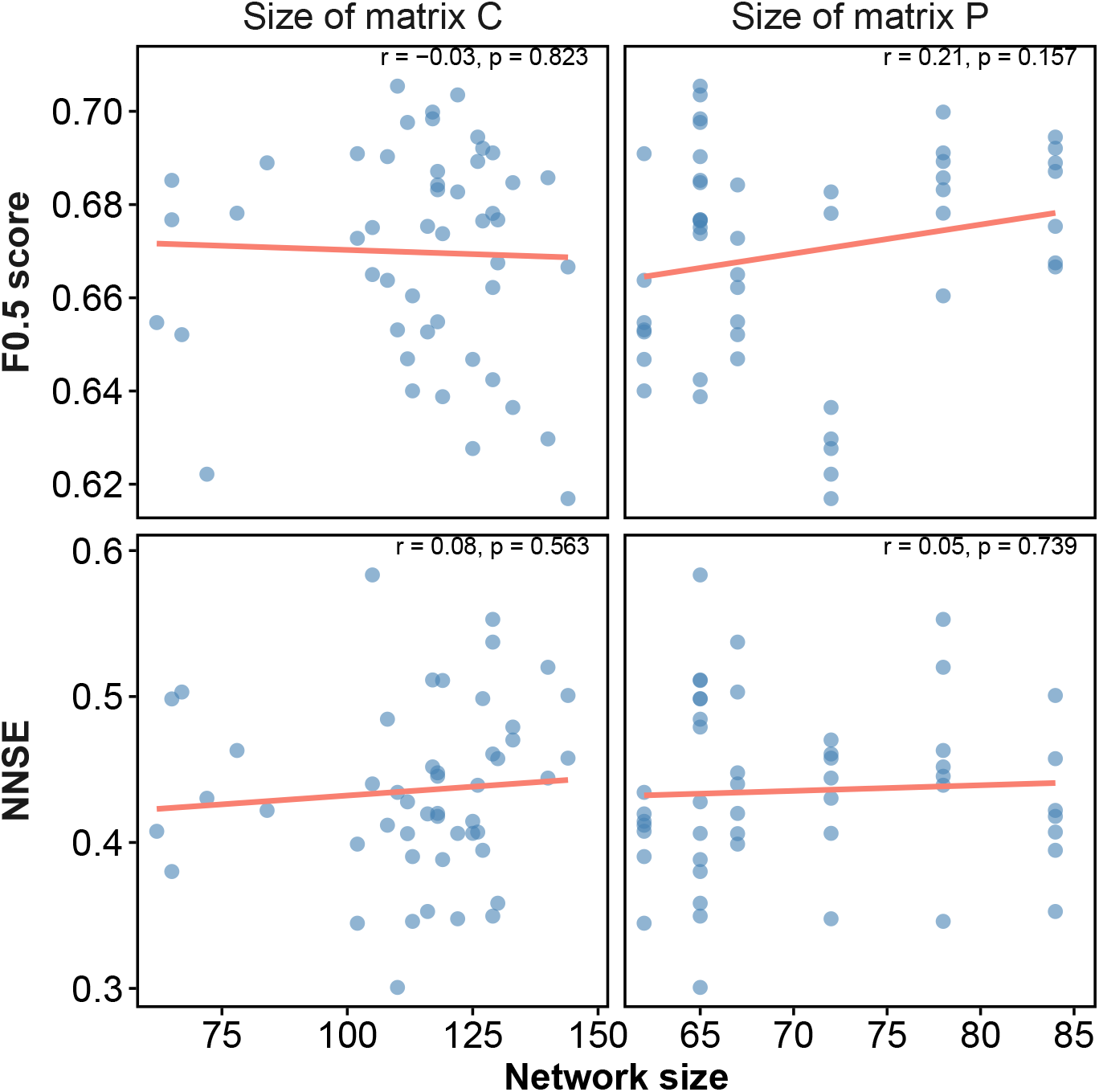
Network size and predictive ability. Prediction quality, for both binary (upper panels) and weighted (lower panels) networks, does not improve with increasing network size.

## 9.3 Appendix S3 Host-parasite network analysis

**Fig. S20:**
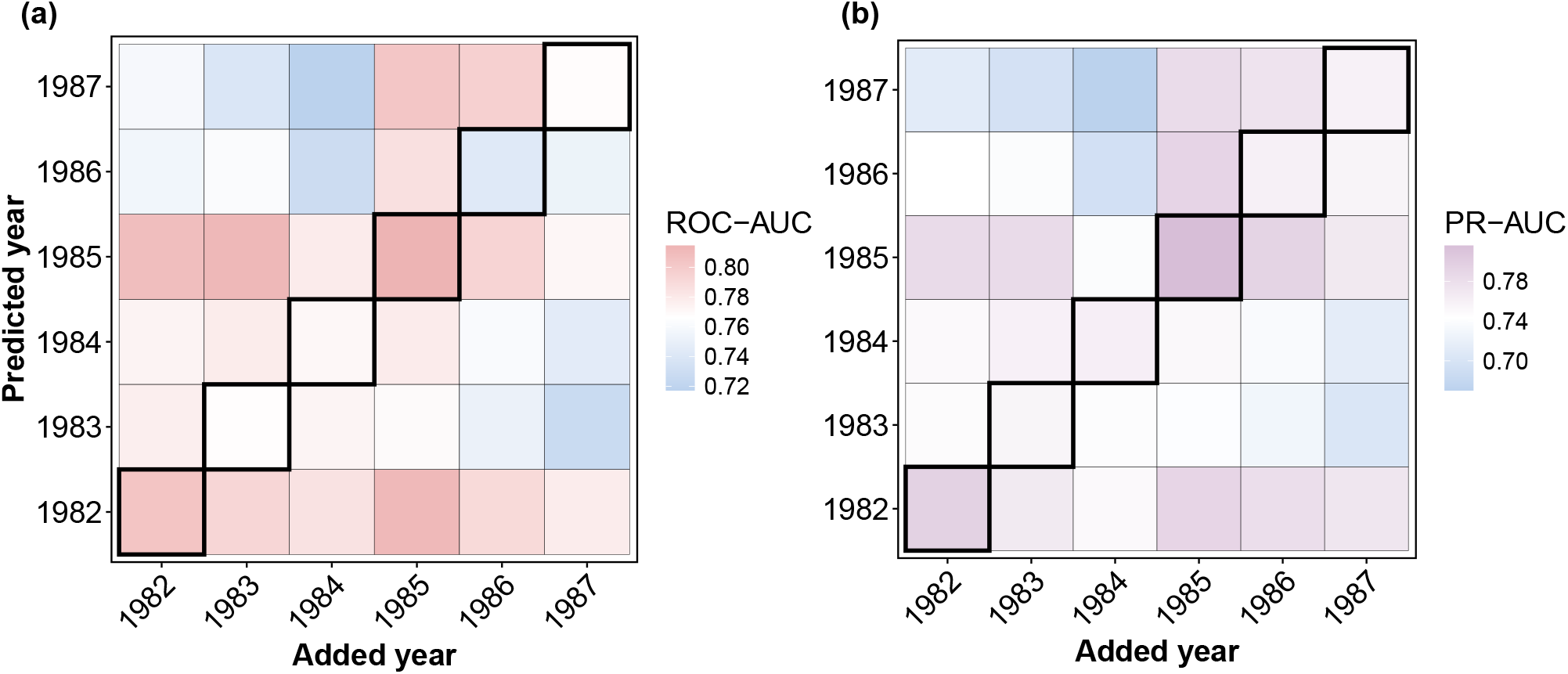
Non-thresholded binary predictive performance for a temporal host-parasite network. **(a)** ROC-AUC values and **(b)** PR-AUC values vary according to the combination of predicted and added networks from different years.

**Fig. S21:**
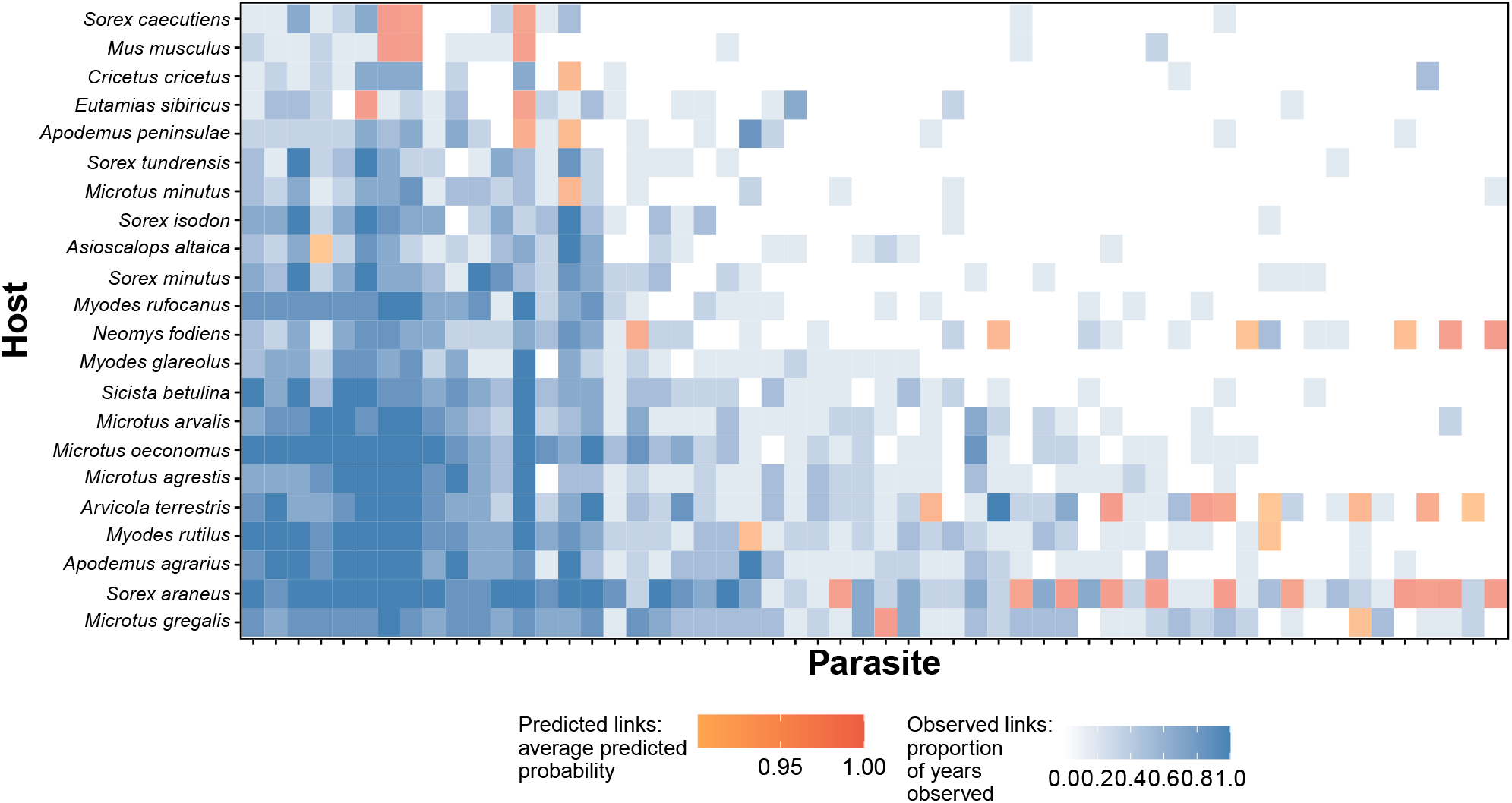
Mapping potential missing interactions in a temporal host-parasite network. **(a)** Blue cells represent observed interactions; darker shades indicate a higher proportion of years in which the interaction was recorded. Orange cells indicate unobserved interactions (interactions never observed in the data), that were predicted to exist based on an average predicted probability above the classification threshold (here, 0.9) across withholding instances within 36 prediction cases (including auxiliary and self-predictions). The intensity of orange reflects this average predicted probability. Species are ordered according to their overall observed degree (total number of unique interaction partners across years). The degree dependence in link assignment, evident in the pollination network (Fig. 3 in the main text) was not observed when applying the same pipeline to this denser host-parasite network.

**Fig. S22:**
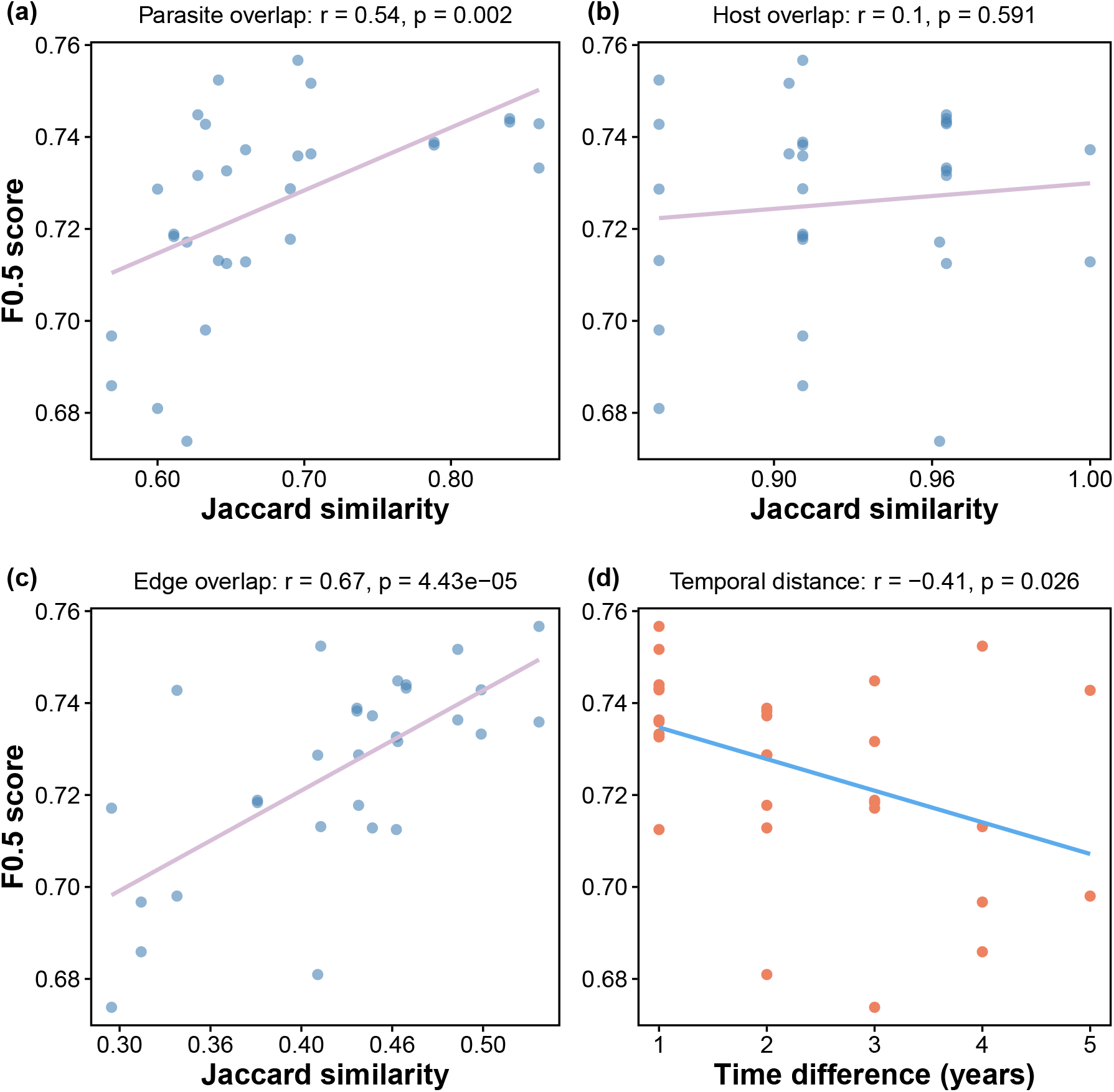
Temporal distance decay in predictive performance. Predictive ability (*F*_0.5_ score) increases with greater similarity in parasite composition **(a)**, shows no relationship with similarity in host composition **(b)**, and increases with interaction overlap between the predicted and auxiliary networks **(c). (d)** Predictive accuracy declines with increasing temporal distance between networks.

